# Evaluating and Improving SSU rRNA PCR Primer Coverage for Bacteria, Archaea, and Eukaryotes Using Metagenomes from Global Ocean Surveys

**DOI:** 10.1101/2020.11.09.375543

**Authors:** Jesse McNichol, Paul M. Berube, Steven J. Biller, Jed A. Fuhrman

## Abstract

Small subunit ribosomal RNA (SSU rRNA) amplicon sequencing can quantitatively and comprehensively profile natural microbiomes, representing a critically important tool for studying diverse global ecosystems. However, results will only be accurate if PCR primers perfectly match the rRNA of all organisms present. To evaluate how well marine microorganisms across all 3 domains are detected by this method, we compared commonly-used primers with > 300 million rRNA gene sequences retrieved from globally-distributed marine metagenomes. The best-performing primers when comparing to 16S rRNA of Bacteria and Archaea were 515Y/926R and 515Y/806RB, which perfectly matched over 96% of all sequences. Considering Cyanobacteria and Chloroplast 16S rRNA, 515Y/926R had the highest coverage (99%), making this set ideal for quantifying marine primary producers. For eukaryotic 18S rRNA sequences, 515Y/926R also performed best (88%), followed by V4R/V4RB (18S rRNA-specific; 82%) – demonstrating that the 515Y/926R combination performs best overall for all 3 domains. Using Atlantic and Pacific Ocean samples, we demonstrate high correspondence between 515Y/926R amplicon abundances (generated for this study) and metagenomic 16S rRNA (median R^2^=0.98, n=272), indicating amplicons can produce equally accurate community composition data versus shotgun metagenomics. Our analysis also revealed that expected performance of all primer sets could be improved with minor modifications, pointing toward a nearly-completely universal primer set that could accurately quantify biogeochemically-important taxa in ecosystems ranging from the deep-sea to the surface. In addition, our reproducible bioinformatic workflow can guide microbiome researchers studying different ecosystems or human health to similarly improve existing primers and generate more accurate quantitative amplicon data.

**Significance Statement:** PCR amplification and sequencing of marker genes is a low-cost technique for monitoring prokaryotic and eukaryotic microbial communities across space and time, but will only work optimally if environmental organisms match PCR primer sequences exactly. In this study, we evaluated how well primers match globally-distributed short-read oceanic metagenomes. Our results demonstrate primer sets vary widely in performance, and that at least for marine systems, rRNA amplicon data from some primers lack significant biases compared to metagenomes. We also show that it is possible to create a nearly universal primer set for diverse saline environments by defining a specific mixture of a few dozen oligonucleotides and present a software pipeline that can guide rational design of primers for any environment with available meta’omic data.

## Introduction

Amplicon sequencing is a powerful tool for understanding microbial community composition and dynamics in the oceans and other ecosystems (1), but the PCR amplification step is potentially biased due to both technical issues during amplification and mismatches to organisms found in natural environments (2–6). Despite these concerns, PCR amplicon sequencing retains several advantages that make it desirable to investigate and correct biases. Firstly, it is a high-throughput and low-cost technique, making it suitable for large numbers of samples - e.g. for global surveys of sediment, water, animal-associated, and other microbial communities (1).

Secondly, the targeted nature of the PCR assay means relatively small numbers of sequences are sufficient for detecting rare organisms even when we have no genomes from any of their relatives, due to the conserved nature of the molecule and the existence of comprehensive SSU rRNA sequence databases. Being able to quantify the abundance and dynamics of rare organisms is important for understanding ecosystem function since many rare bacteria have impacts far greater than their low abundances might imply (7, 8). While PCR assays are targeted, we note that this does not imply they need to be taxonomically-restricted. For example, there are some primer binding sites in the SSU rRNA molecule that are nearly universally conserved between Archaea, Bacteria, and Eukarya (discussed further below).

Thirdly, using untargeted metagenomic sequencing for taxonomic profiling still has a number of significant disadvantages versus amplicon sequencing. Direct taxonomic assignment of randomly- sheared metagenomic reads by recruitment to reference genomes has the potential to assign taxonomy at a very high resolution. However, the best currently available genome databases still only recruit ∼ 40 % of metagenomic reads for marine prokaryotes (9), meaning a significant fraction of untargeted metagenomic datasets are taxonomically uncharted. This problem is even more acute for environmental eukaryotes with large amounts of non-coding genomic DNA and fewer genomic references. It is also possible to extract SSU rRNA (or other marker genes) from metagenomes to generate a taxonomic profile. However, SSU rRNA fragments typically represent only a small fraction a given dataset (∼ 0.1 %), so is a far more costly way to obtain a comprehensive community profile. In addition, because metagenomically-retrieved marker genes are randomly sheared fragments covering both conserved and hypervariable regions and are in most cases too short to provide a unique match to reference sequences, they must be assembled or otherwise clustered using non-deterministic algorithms. Because of this, the taxonomic resolution of short-read metagenomic marker gene data is limited (e.g. near family or genus level for the best known conserved markers like rRNA). In contrast, with rRNA amplicons, modern “denoising” algorithms produce exact amplicon sequence variants (ASVs) that are stable biogeographic markers that can be intercompared without reanalysis (10). The resulting datasets are also relatively simple to analyze since they consist of a single gene region and can be comprehensively classified with databases such as SILVA or RDP (11, 12).

Recent studies have shown that PCR amplicon sequencing of mock microbial communities can recover known relative and absolute abundances extremely well, and thus provides accurate quantification of natural gene copy abundances (3, 13, 14). In turn, this accuracy allows amplicon data to serve as a ground-truth for modeling / ecological studies by quantifying community members and their dynamics. While well-designed primers have the potential to recover truly quantitative data, these studies also underscored the critical fact that a single mismatch between the primer and template sequences can have a dramatic effect on the measured community composition in complex natural mixtures. For example, a single internal mismatch in the popular Earth Microbiome Project primer caused a ∼10-fold bias against the most common bacteria in seawater (SAR11 cluster), and terminal 3’ mismatches can completely prevent amplification (2, 3, 6, 13). Since the extent of this bias is not easily predicted, PCR primers should incorporate degenerate bases or specific oligonucleotide variants so that all targeted organisms are perfectly matched without overly diluting the common perfect matches in primer mixtures. This is especially critical for abundant taxa such as SAR11 as distortions in their relative abundances will skew the remainder of the community, but it is also important to consider for rare taxa which might have essential biogeochemical or ecological roles.

Previous studies have shown high coverage of natural taxa is possible by designing moderately degenerate primers without sacrificing specificity or PCR efficiency (3, 13). This primer design was accomplished by comparing oligonucleotide sequences to a SSU rRNA database such as SILVA or RDP (11, 12) and then checking for mismatches to organisms known to be abundant in the environment of interest. This approach led to marked improvements in primer design, for example by Apprill et al (2015) and Parada et al (2015) who reported that small modifications to existing primers could better quantify the dominant marine taxa SAR11 and *Thaumarchaea* (2, 3). However, in these primer evaluations, the reliance on full-length references and giving each sequence in a database equal weight can lead to a distorted perspective of the actual extent of matches and mismatches expected in real samples since they do not take into account the highly unequal abundances in nature. In addition, some environments may have abundant taxa poorly represented or unrepresented in these curated reference databases.

Wear et al. (2018) studied the effect of these potential biases empirically by testing 4 primer sets currently in broad use by marine microbiologists on a 16S rRNA mock microbial community derived from natural seawater communities near Santa Barbara (6). These authors tested the effect of their analysis pipeline on recovery of mock community sequences for different primers, and found their primer-pipeline combination had several sequence-specific biases, in some cases due to a primer mismatch. While an important step towards cross-comparing primers in an oceanographic context, Wear et al. note that results are specific to their mock microbial community, analyzed with their particular pipeline, and thus it remains unknown how representative their results are for other ecosystems and other pipelines. More specifically, it is currently unknown how many primer- mismatched rRNA sequences occur across diverse oceanographic environments, a key piece of information for designing field measurements and interpreting results.

To more fully evaluate the extent to which primers currently in broad use by the marine microbiology community perfectly match naturally occurring sequences, we developed a workflow to conduct *in silico* primer evaluations with metagenomic SSU rRNA fragments, under the general assumption that metagenomes have fewer methodological biases than PCR. While it is already known that some primers have better coverage for abundant marine organisms versus others (3, 13), a precise quantitative comparison of these primers with others that are in broad use has not yet been conducted.

We developed a new software pipeline for analyzing globally-distributed marine metagenomic datasets (Table S1) to make a quantitative and objective evaluation of PCR primers for pelagic marine ecosystems and to suggest specific improvements based on this evidence. We further experimentally compared the quantitative performance of SSU rRNA amplicon sequencing, performed for this study, against the published shotgun metagenomic data from the same sample DNA from the bioGEOTRACES study. Our pipeline also allowed us to make intercomparisons between primer sets often not considered together. For example, we compared the performance of two “universal” (3- domain) PCR primers that amplify both 16S and 18S rRNA with primers designed to specifically target only 16S or 18S rRNA and exclude the other molecule.

Because our software pipeline is broadly applicable for any environment where metagenomic data exists, it will allow investigators to design environment-specific PCR probes that recover as many (or as few) targeted organisms as desired. It is also flexible, allowing design or improvement of primers from different variable regions to either retain stable intercomparisons with existing data, or to target a region of the SSU rRNA molecule that has better taxonomic resolution for taxa of interest (15). Our approach has the added advantage of providing a quantitative, evidence-based framework to identify which taxa may have been affected by biases in existing datasets generated with sub-optimal primers - information that is often unknown or only recognized anecdotally by specific investigators.

## Results and Discussion

### Overall primer coverage for pelagic ocean datasets

To evaluate the real-world extent of primer biases, we compared PCR primers currently in broad use by the marine microbiology community (Table S2) to SSU rRNA sequences extracted from a geographically diverse set of pelagic ocean metagenomes (Fig 1; Table S1). These analyses were based on the percentage of perfect matches between primers and metagenome sequences as a metric to determine potential PCR performance, since previous studies have shown that even a single mismatch can significantly bias results (3, 13).

**Figure 1:**
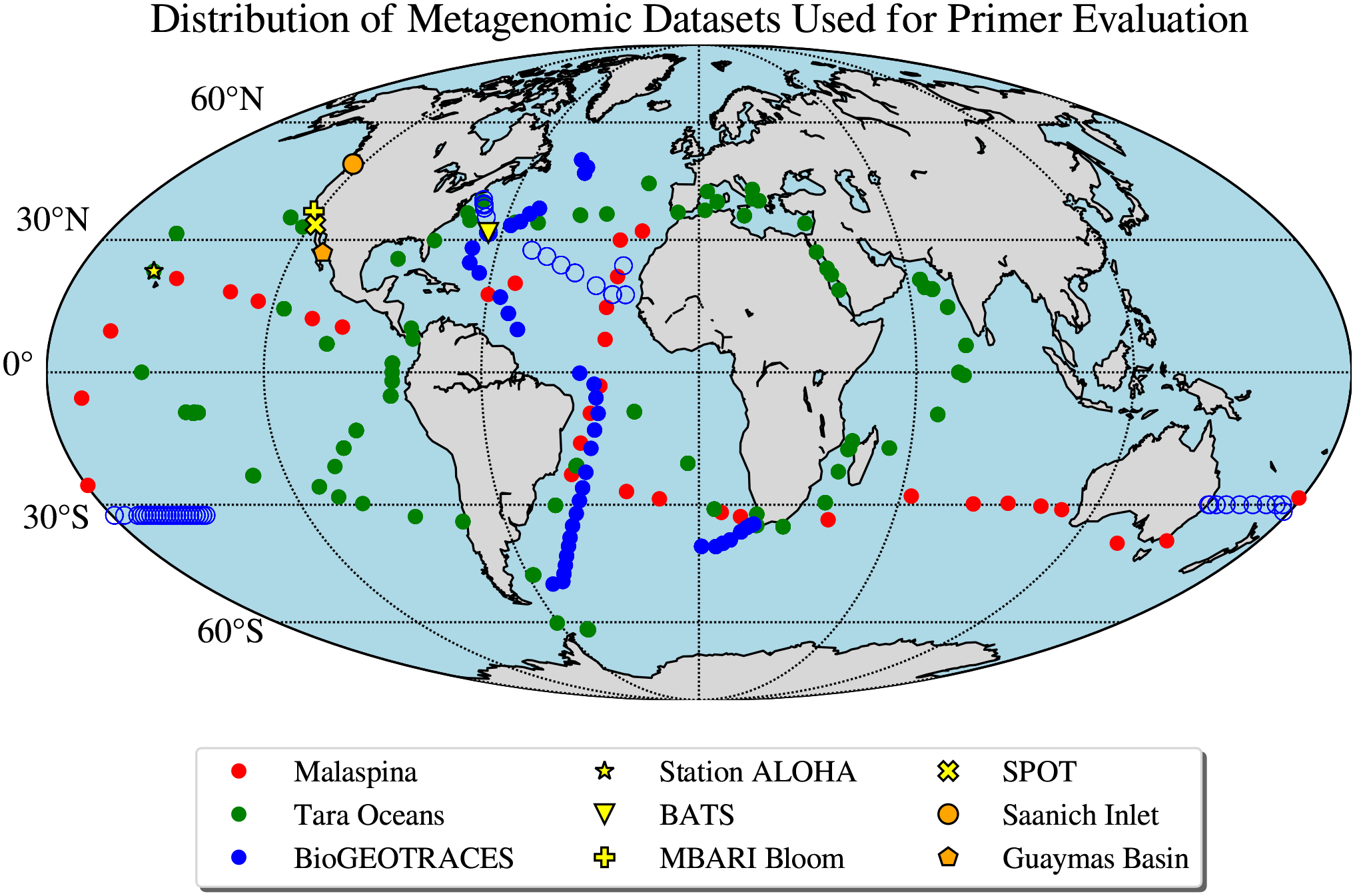
Distribution of metagenomic samples used in this study. BioGEOTRACES samples that were used in the metagenome/amplicon intercomparison are noted with open circles (GA03 - N. Atlantic; GP13 - S. Pacific).

We observed relatively even recruitment of metagenomic reads across the SSU rRNA molecule, which indicates that each different primer region has broadly similar potential to generate accurate taxonomic profiles of naturally-occurring organisms (Supplemental Results). However, when we compared the precise sequences derived from the primer-binding regions to the oligonucleotide primer sequences, we observed that predicted median coverage of perfect matches varied widely among primer pairs (Table S2), with 515Y/806RB and 515Y/926R showing the most consistently high incidence of perfect matches (Fig 2). The ability of all primers to capture the underlying diversity in the metagenomic samples can be improved to some extent by increasing degeneracies, though this varied significantly between primer sets. Adding a single additional degeneracy fixes the majority of mismatches for some primers (e.g. 785R; discussed further below), whereas others have greater than 2 mismatches to metagenomic sequences and thus would require more extensive modifications (Fig S1-S28). Below, we discuss the performance of these primers across 4 broad taxonomic groups (*Archaea, Bacteria, Cyanobacteria+*Plastidal 16S rRNA, and *Eukarya*). We separated *Cyanobacteria* + phytoplankton plastidal 16S rRNA from bacterial 16S rRNA since these organisms are responsible for the vast majority of marine primary productivity and thus are important to quantify accurately.

**Figure 2:**
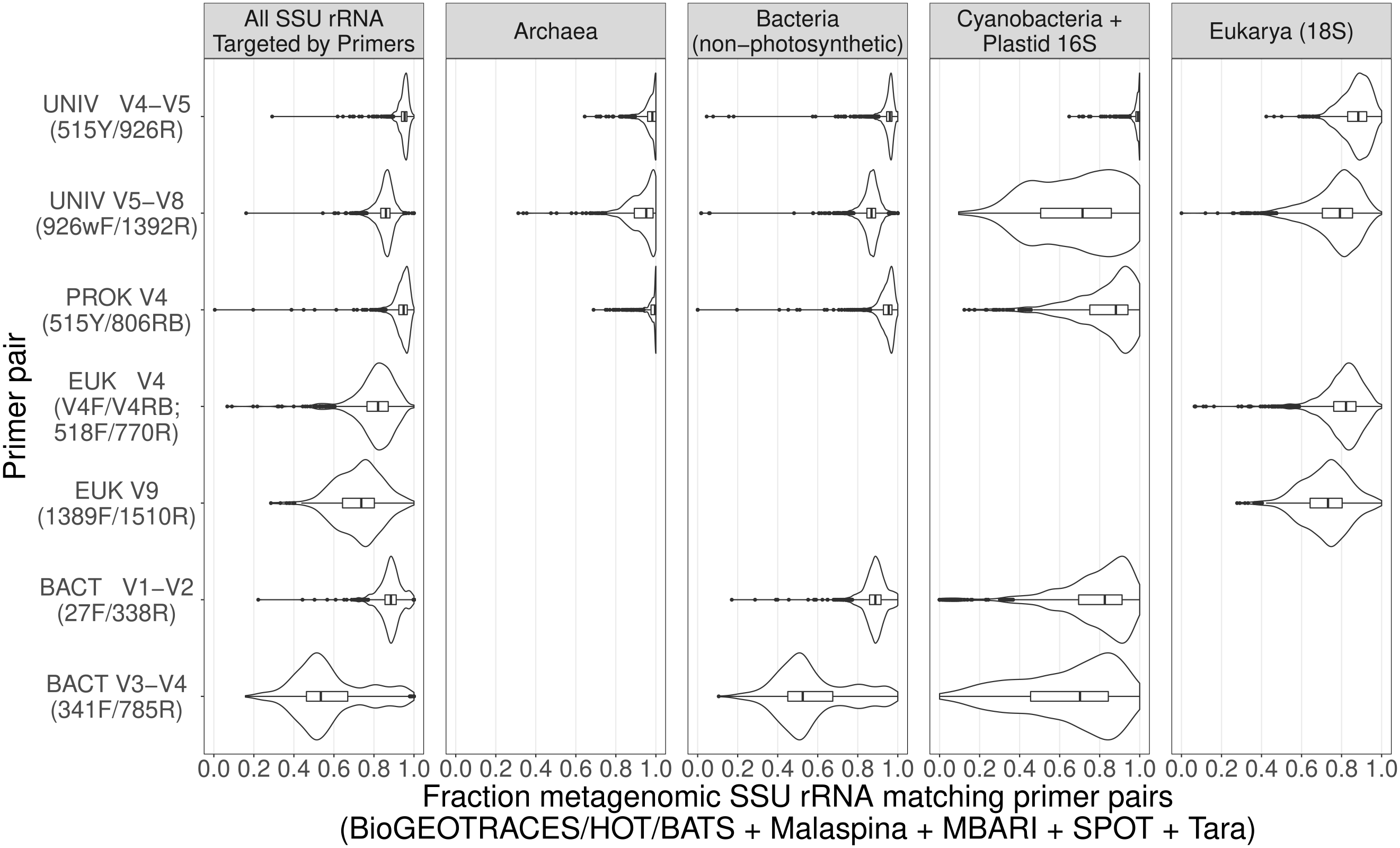
Oligonucleotide PCR primer coverage based on comparisons with globally-distributed oceanic metagenomes. The shapes of the plots illustrate the distribution of primer coverage across all samples, where perfect correspondence of primers and metagenome sequences is represented by data piled up against “1.0” on the x axis, a situation approached by the UNIV V4-V5 primer for Cyanobacteria+ Plastid 16S rRNA, for example. Note that the “All SSU rRNA” panel represents all of the SSU rRNA targeted by a particular primer set (e.g. for 1389F/1510R this includes only Eukarya while for 926wF/1392R it includes all 4 taxonomic groups). Data are corrected for predicted taxonomic overlap between forward and reverse primers. Coordinates refer to locations of primer alignments to the Escherichia coli 16S rRNA gene (strain K-12; substrain MG1655).

For *Archaea*, the primer with the best median coverage was the EMP 515Y/806RB combination (coverage=0.996), followed by 515Y/926R (0.982), and 926wF/1392R (0.952). We also note that our results suggest the 341F/785R primer combination could amplify most *Archaea* with the addition of several degeneracies in 341F (Fig S1-S7).

For *Bacteria* (excluding *Cyanobacteria* and plastidal 16S rRNA sequences), the primer with the best median coverage was the 515Y/926R combination (0.961), followed by 515Y/806RB (0.955), 27F/338R (0.888), 926wF/1392R (0.869), and 341F/785R (0.525). The relatively poor performance of 341F/785R is mainly due to the reverse primer (median individual primer coverages: 341F=0.887, 785R=0.624), whereas the other two less optimal primer pairs have more even individual performances (27F=0.968, 338R=0.913; 926wF=0.916, 1392R=0.943).

For *Cyanobacteria* and chloroplast 16S rRNA (derived from eukaryotic phytoplankton), the primer pair with the highest median coverage was 515Y/926R (0.992), followed by 515Y/806RB (0.881), 27F/338R (0.826), 926wF/1392R (0.714), and 341F/785R (0.702). We note that the vast majority of these mismatched sequences come from chloroplast 16S rRNA sequences - the common cyanobacterial groups *Prochlorococcus* and *Synechococcus* have high coverage with all primers listed above, so this is only a concern for those wishing to use chloroplast 16S rRNA data to quantify eukaryotic phytoplankton.

For *Eukarya*, the primer pair with the highest median coverage was 515Y/926R (0.883), followed by V4F/V4RB (0.822), 926wF/1392R (0.792), and 1389F/1510R (0.732). The distributions of coverage were considerably broader for *Eukarya* versus all other 16S rRNA categories, which may be due to a higher sequence heterogeneity among eukaryotic genomes. As above, all primer sets could be improved with the addition of some degeneracy (Fig S22-S28), but approaching perfect coverage seems less practically achievable for eukaryotic SSU rRNA sequences. In addition, our analysis indicates that improvements in eukaryotic coverage are most likely to come from targeting universally- conserved rRNA regions, such as those containing the 515Y/V4F and the 1389F/1392R primers respectively (Fig S22-S28).

We note, however, that recognizable dinoflagellate chloroplast rRNA sequences were almost completely missing from the metagenomic sequences (for all primer regions, not just 515Y/926R), and thus may not appear in the resulting amplicons regardless of primer choice. This is possibly due to the unusual genomic organization/subcellular localization of chloroplast genes in dinoflagellates (16), or alternatively because we lack insufficiently accurate or comprehensive databases for identifying dinoflagellate chloroplast 16S rRNA (discussed further below). It is possible, however, to recover and confidently identify dinoflagellate 18S rRNA sequences from the 515Y/926R primer pair (17), making it feasible to reconstruct a holistic picture of phytoplankton community composition / abundance in combination with the chloroplast 16S rRNA.

In summary, the primer pairs with the highest environmental coverage were 515Y/926R (universal), 515Y/806RB (prokaryote-only), and V4F/V4RB (eukaryote-only). It is worth emphasizing that both universal primer sets have the potential to be equally or more comprehensive for *Eukarya* versus the *Eukarya*-specific primer sets tested here. In addition, for those wishing to quantify the abundance of eukaryotic phytoplankton using chloroplast data, the 515Y/926R primer set nearly perfectly matches all environmental sequences. These results are a quantitative confirmation that the careful primer design previously reported for the 515Y/806RB/926R primers has resulted in very high overall coverage in oceanic ecosystems. On the other hand, certain primer sets (e.g. 341F/785R) have very low coverage due to mismatches to dominant organisms, despite being designed to maximize environmental coverage (18). Resulting data may thus be distorted, though we note that removing taxa with mismatches to primers from processed data will recover accurate relative abundances for the remaining taxa (13), due to the fact that biases are thought to be taxon-specific (19). This relative abundance correction approach depends on our pipeline’s specific identification of mismatched taxa, and provides a way to recover useful quantitative information from legacy datasets that may have been biased during PCR. In combination with ecosystem-specific full-length 16S rRNA databases, this correction approach could represent an important tool for integrating historical amplicon datasets and modern long-read sequencing data into a single intercomparable framework for observing longer-term changes in ecosystem structure (15). In addition, by eliminating or at least controlling for primer bias, our approach could help better evaluate how well particular primer regions perform with respect to taxonomic profiling and classification since it would tease apart the effect of variable region from amplification artifacts.

### A Quantitative Intercomparison between SSU rRNA Amplicon Sequencing and Metagenomics

In order for amplicon sequencing to become useful as a quantitative tool for measuring natural gene abundances, it is desirable to benchmark amplicon abundances against an external reference that is not affected by potential PCR biases to gain a more objective determination of how well a primer recovers true gene abundances. While shotgun metagenomics is not necessarily free of all biases, these biases are distinct from those in PCR-based amplicon studies; thus, if both data types yield similar community composition patterns, we can be more confident about our overall conclusions. We tested whether amplicon-based strategies recover similar patterns as metagenomes by generating PCR amplicon sequences for a subset of 272 BioGEOTRACES (20) samples (cruises GA03 and GP13). We used the same DNA, eliminating extraction biases as a potential confounding factor in this intercomparison. We were thus able to make a direct “apples to apples” quantitative comparison between the relative abundances of organisms determined with amplicons vs. metagenomic reads.

To accomplish this, we compared exact amplicon sequence variants (ASVs) from the 515Y/926R primer pair with metagenomic reads from the same region of the SSU rRNA molecule. To make a more robust quantitative comparison, ASVs were clustered as necessary to account for the fact that short (150bp) metagenomic reads could in some cases only be attributed to a broader phylogenetic group (e.g. *Prochlorococcus* and SAR11). The summed abundance of all taxonomic groups, as determined by ASVs and metagenomic reads, was then compared using a linear least-squares regression. For 16S rRNA sequences, we observed very similar community composition for both techniques, as shown by very high R^2^ values and coverage in Fig 3a (median R^2^=0.98; range=0.55-1.0). This indicated that the ASV sequences were recovering the same diversity of organisms present in metagenomic reads from the primer region and is consistent with the aforementioned observation of even coverage of metagenomic reads across the SSU rRNA molecule.

**Fig 3:**
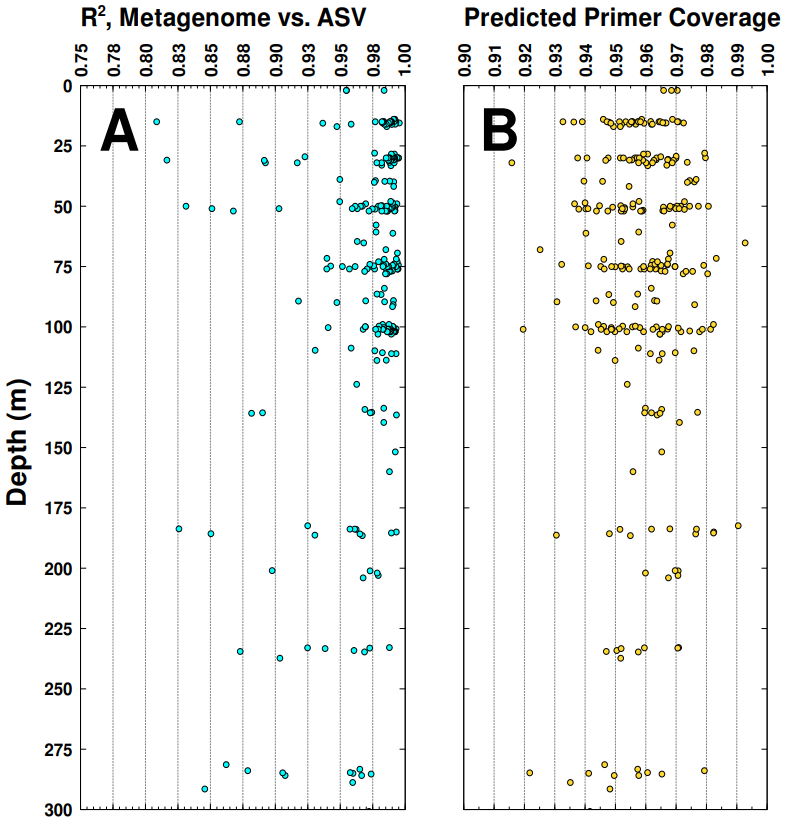
Metagenome-amplicon/oligonucleotide primer comparisons across the marine water column. A) Metagenome-amplicon quantitative correspondence across all depths for the GA03 cruise (BioGEOTRACES), showing the R2 value of the plot of the relative abundances of amplicon sequence variants (ASVs; 515Y/926R primers) vs. relative abundances of metagenomic reads from the same organisms (see text for details of comparisons). B) The fraction of exact matches of the 515Y and 926R amplicon primers compared to metagenomic reads of the priming regions for each sample, summarized as the “worst-case” coverage for the combined primers (i.e. 1 - (missed_fwd_ + missed_rev_)).

These patterns were generally consistent across the surface-mesopelagic transition where there is a major change in community composition (Fig 3). We do however note that at depths > 150m about 38% of samples had R^2^ values below 0.95 (Fig 3a), vs. 9.2% for samples at depths < 150m. These lower correspondences are unlikely to be due to mismatches to the primer region, since predicted primer coverage is uniformly high across depth for the 515Y/926R primer pair across depth (Fig 3b). Other factors such as DNA quality or quantity may have been responsible for this, but we cannot rule out the possibility it involves characteristics of the sequences between the priming regions such as secondary structure or expanded loops known from some eukaryotes (21). Regardless, these results show that amplicon sequencing and metagenomic profiling typically have a very high quantitative correspondence with the primers tested here (515Y/926R). Additionally we provide a software toolbox for making further intercomparisons between arbitrary primer sets and metagenomes that could be useful for optimizing both PCR amplicon sequencing and metagenomics for diverse environments.

### Improving Existing Primers

In addition to quantifying mismatches, our pipeline also identifies the specific primer region variants found in particular naturally-occurring taxa and their coverage across datasets. This information can guide improvements for any of the primers tested here, and we identify two specific use cases. Firstly, some of the primers identified as having major mismatches are dominated by a few organisms, and thus small modifications are all that is necessary to make them more appropriate for oceanic ecosystems. Secondly, we were interested to know whether for the better-performing primers it would be possible to identify a combination of oligonucleotides that would perform well across all marine environments tested here. Such a primer set could have the advantage of being able to monitor rare but biogeochemically-important taxa that may become more abundant due to climate change, e.g. as a result of expanding oxygen minimum zones.

To illustrate how a primer with poor predicted performance could be improved, we use the case of 785R which has a mismatch to the dominant organism SAR11. Because the SAR11 mismatches represent up to 40% of total 16S rRNA reads (Table S3), this one mismatch will have a major effect on the resulting data but could be corrected by adding a single additional degeneracy at position 7 in the primer (5’-GACTAC**<u>N</u>**VGGGTATCTAATCC). However, 785R also has mismatches to a number of abundant chloroplast sequences (Fig 2), which could be corrected with two additional degeneracies (5’- **<u>R</u>**A**<u>Y</u>**TACNVGGGTATCTAATCC). Compared to the original primer (5’- GACTACHVGGGTATCTAATCC), these primers would require the synthesis of a larger number of oligonucleotide combinations (12 and 48 vs. 9 originally) but are well within the number of combinations currently in use for other environmental amplification strategies (22). Interestingly, despite being the worst-performing primer for marine ecosystems due to mismatches to the dominant taxon SAR11, we predict that by adding the above degeneracies this primer would achieve nearly perfect coverage for *Bacteria* (Fig S15-S21). This suggests the primer-binding region of 785R is highly-conserved among *Bacteria,* consistent with the initial publication describing this primer as the best overall for prokaryotic taxa across diverse environments (18). This observation also underscores the value of comparing primer sequences with environmental data since it uncovered a significant, unexpected mismatch to an abundant taxon that can now be corrected.

We also investigated whether it is possible to improve primer sets that already perform well (e.g. 515Y/806RB/926R) to the point where they offer near-universal coverage of all taxa identified in this study. To do so, we identified datasets that had significant potential for coverage improvement (Figures S1-S28) and certain rare order-level taxa that had many mismatches or are known to be biogeochemically-important.

Our data showed that the 806RB primer performed extremely well overall for 16S rRNA sequences, but did miss a number of 16S rRNA chloroplast sequences mostly from the *Mamielliales* clade, which includes small picoeukaryotes such as *Ostreococcus* and *Bathycoccus*. As shown in Fig S8-S14, these taxa were still missed when allowing for up to 2 mismatches, meaning they are unlikely to be accurately quantified with the current primer design. Alterations would be possible, but we note that this may be an intentional design choice, since chloroplast amplification may be undesirable in certain environments (23) but is often very useful in euphotic zone marine water samples where it can be used to quantify primary producer abundance. Related to this, we note that a context-dependent advantage of 806RB is that it lacks an 18S rRNA binding site. This means that in combination with 515Y, nuclear 18S rRNA will not be amplified with 806RB. In other words, even though the variable region covered by 515Y/806RB contains eukaryotic sequences, this primer set will not recover them as amplicons – demonstrating that primer design places a first-order constraint on how well a given amplicon product reflects the natural community. In contrast, the 51Y/926R combination will amplify these eukaryotic sequences. In a practical sense, amplifying 18S rRNA sequences with 16S rRNA can present problems, for example in host-associated microbiome studies where 18S rRNA could overwhelm targeted 16S rRNA. This overwhelming of libraries by 18S rRNA sequences is however unlikely to be a major issue for the 515Y/926R primer pair for pelagic ocean samples. This is due to the fact that 18S rRNA sequences will generally be selected against in the sequencing process (thus reducing their overall abundances (13)), and the fact that marine environments are generally dominated by prokaryotic microorganisms (13; Fig S29).

The 515Y primer was relatively easy to improve, given that its performance was already very robust across diverse datasets (Fig S1-S28). We identified four positions where further degeneracies could be added (5’-GTG**<u>B</u>**CAGCM**<u>SY</u>**CGCGGT**<u>M</u>**A), and were sufficient to resolve the vast majority of mismatches including to rare taxa such as certain representatives of the *Patescibacteria* (a.k.a. candidate phyla radiation (CPR) bacteria) which were previously identified as taxonomic “blind spots” in PCR amplicon analysis (5). However, incorporating these new degeneracies produced a relatively large number of oligonucleotide combinations compared to the original primer (48 versus 4 originally) and most of these new variants were not observed in the metagenomic data. We thus decided to redesign the primer based on a different approach, starting from a non-degenerate primer and only adding in variants that could be detected in the natural environment or in the SILVA database. This resulted in a specific mixture of 13 specific oligonucleotides that we term 515Yp-min (“p” for pelagic, “min” for minimal). This mixture of oligonucleotides is tailored specifically to the environmental sequences present in natural marine environments. It both maximizes organismal coverage, while keeping the overall degeneracy at a relatively low level compared to other degenerate primers used in environmental microbiology (22). Another key advantage of this tailored approach is that it permits iterative improvements to a primer set over time. For example, we added a variant matching dinoflagellate chloroplast sequences from the PhytoRef database (24) that may potentially improve quantification of these organisms’ plastidal 16S rRNA in combination with other modifications to 926R (discussed below). In the future, if we were able to identify other variants explaining why dinoflagellate chloroplast 16S sequences are conspicuously absent from 515Y/926R amplicon data, it would then be feasible to add these new olignucleotides to the mixture without creating excessive sequence redundancy.

The 926R primer had mismatches to more taxa, and thus required more extensive modifications. For example, Table S3 shows that 926R has mismatches to both *Ectothiorhodospirales* (an uncultivated group related to purple sulfur bacteria (25)), and the *Rickettsiales* (an order that includes mitochondrial sequences in addition to free-living organisms (26)). While less abundant overall, we also noted mismatches in the Tara Oceans dataset to *Brocadiales*, the order containing anammox bacteria (27). In the Guaymas basin sediment samples, some of the dominant mismatches were to the *Campylobacteria,* dominant chemoautotrophs in many sulfidic systems (28), as well as unusual *Archaea* that are now recognized to be much more diverse than previously thought and missed by some PCR primers (29). Accurate quantification of these rare taxa would allow the monitoring of unusual but potentially significant changes in biogeochemistry. For example, the *Campylobacteria* are known to be abundant in certain low-oxygen microenvironments such as in sediment trap particles (30), and the *Brocadiales* are abundant in oxygen minimum zones (27). Being able to better quantify these “sentinel taxa” could allow us to monitor the dynamics of these low-oxygen environments that are likely to expand under global climate change.

Following the same approach discussed above for 515Yp-min, we developed a mixture of 38 oligonucleotides (“926Rp-min”) that contains perfect matches for aforementioned taxa, primer variants that have > 2% overall relative abundance across all environments, as well as 2 sequences from dinoflagellate chloroplasts that were not previously covered. This set of oligonucleotides is comprehensive for all the environments tested in our study and would likely result in major improvements in quantification all of the aforementioned taxa. To achieve the same organismal coverage by adding more degenerate sites would increase the total number of primer variants to at least 144 (compared to 16 in the original primer). Not only would this dilute perfectly-matching oligonucleotides and be unlikely to result in efficient PCR amplification, it would also include many redundant oligonucleotides unnecessary with an explicit oligonucleotide mixture. The improvements in coverage for this new mixture may be particularly apparent for *Rickettsiales* templates, many of which have a single mismatch at the 3’ end of the primer that is known to completely prevent PCR amplification (6). Since the order *Rickettsiales* contains sequences from mitochondrial SSU rRNA, this modification will potentially allow for more effective quantification of protist abundances, similar to how chloroplasts quantify marine protistan phytoplankton.

Since it is beyond the scope of this paper to provide exhaustive recommendations for each primer set, we direct interested readers to an Open Science Foundation repository which contains all necessary output files, scripts, and a suggested workflow for improving primers using metagenomic data (osf.io/gr4nc/). Improvements should be possible for most taxa/primer combinations since a few mismatches typically dominated, with a long tail of rare variants that may derive from sequencing errors or pseudogenized SSU rRNA. This information would allow an interested reader to optimize one of the primers investigated here for particular taxa or environments of interest.

Moving forward, we recommend oceanographers consider applying these two new primer mixtures for pelagic water column surveys to maximize organismal coverage (515Yp-min, 926Rp-min; Table S4). By correcting for 3’ mismatches that lead to taxonomic “blind spots” and including exact matches to taxa found in low-oxygen regions, we should be able to better quantify the abundance and dynamics of organisms critical to marine carbon, nitrogen, and sulfur cycling. In addition, by representing primers as a specific mixture of oligonucleotides rather than a single sequence with degeneracies, it leaves open the possibility of small modifications in the future to improve coverage; the same is not true for fully degenerate primers where each new degeneracy will multiply the total number of combinations by at least 2-fold and further dilute the other perfectly matching oligonucleotides. If proven to be as quantitative as the original 515Y/926R primers (3, 13), these new mixtures would provide an affordable, comprehensive (all organisms from Archaea - to Zooplankton), and scalable method to measure whole community compositions across time and space in the world’s oceans. While further validation would be necessary, we believe our results and those of previous studies (3, 13–15) demonstrate the potential for PCR amplicon methods to accurately quantify natural marker gene abundances in a robust and accurate manner, either by accurately recovering true relative abundances, or in combination with internal standards to obtain absolute quantification (14). This would allow confident measurement of microbial community composition alongside well-developed methods such as those for quantifying phytoplankton photosynthetic pigments. In turn, this would help expand our understanding of how oceanic microbes interact with biogeochemical cycles and respond to global climate change.

Finally, we believe our bioinformatic approach could be applied elsewhere to generate ecosystem-specific primer sets. For example, it remains unknown whether the patterns we observed for seawater would be true in other systems, such as terrestrial sediment, soil, freshwater, or animal/plant- associated microbiomes. Our reproducible workflow, combined with expert curation and broad environmental sampling could ensure that taxa of interest are accurately quantified by oligonucleotide primers for any environment where deeply-sampled metagenomes could be used as a database. In addition, our approach could allow for the rational evaluation and improvement of new primers for next generation long-read amplicon sequencing, or to improve the performance of primers used to identify animal species of economic or conservation importance. By evaluating primer coverage objectively and providing simple ways to iteratively improve existing primer sets, it would ensure that the potential of these techniques for monitoring microbiomes relevant to ecosystem functioning and human health is fully realized.

## Materials and Methods

### Sample scope / biogeographic distribution and evaluated primers

We analyzed globally-distributed metagenomic datasets that reflect a range of depths and nutrient regimes (Table S1; Fig 1), and tested primers that are commonly used in environmental microbiology (Table S2; (2–4, 18, 31–35)). Metagenomic samples included worldwide sampling expeditions such as Tara oceans (36), the Malaspina expedition (37), and BioGEOTRACES (20). Among these three campaigns, the Malaspina metagenomes focus largely on the deep sea, whereas TARA and BioGEOTRACES sample primarily the sun-lit ocean. In addition, we used time-series data from the Hawaii Ocean Time-Series and the Bermuda Atlantic Time-Series (38, 39) (HOT/BATS; both analyzed with BioGEOTRACES as BiHOBA [BioGEOTRACES-HOT-BATS]), a bloom time-series in Monterey Bay (40), and the San Pedro Ocean Time-Series (41) (SPOT). We also included in the analysis two sites representing suboxic/anoxic systems - the Saanich inlet time-series (42), and samples collected from hydrothermally-altered sediments in Guaymas basin (43). We also generated new 16S/18S rRNA universal amplicon sequences for two BioGEOTRACES cruise transects (GA03/GP13; noted with open circles on Fig 1) which were used for making intercomparisons between metagenomes and amplicons.

### Evaluation of primer coverage/biases using metagenomic reads

In order to evaluate the performance of our primer set compared to others, we used metagenomic reads to estimate the fraction of SSU rRNA fragments that could be successfully targeted by (i.e. perfectly match) each primer set. In brief, our pipeline retrieves SSU rRNA from fastq- formatted shotgun metagenomic samples, removes sequences containing uninformative repeats, sorts into four organismal categories (*Archaea*, *Bacteria, Cyanobacteria* (including plastidal 16S rRNA), and *Eukarya*), and aligns these sequences to a group-specific reference SSU rRNA sequence (e.g. *Saccharomyces cerevisiae* for *Eukarya*). To obtain sequences overlapping with each oligonucleotide primer and to exclude non-primer reads, coordinates provided in the config file were used to extract reads from alignments that overlapped with the primer-binding region plus 5 leading or trailing bases. These reads, which represent SSU rRNA fragments with a primer-binding region, were then quality- filtered and compared with primer sequences (Table 2) at 0, 1 and 2-mismatch thresholds (Fig S1-S28) to identify matched and mismatched sequences. As an additional quality-control step, we excluded any aligned read falling in the primer region that did not have a match to the primer at a 6-mismatch threshold (e.g. misalignments or distantly homologous sequences). Both mismatched and matched reads were then classified using the SILVA 132 database (*Archaea, Bacteria, Eukarya*) and PhytoRef (11, 24) (*Cyanobacteria +* plastidal 16S rRNA). The specific software implementation is described fully in the supplementary material. For those wishing to inspect results further, complete plaintext summaries of output and summary plots are available online at https://osf.io/gr4nc/.

These computational steps were implemented with the snakemake workflow engine (44) to create a documented and reproducible pipeline which is freely available (<u>github.com/jcmcnch/MGPrimerEval</u>). It automatically produces tabular output / summary plots and can be applied to new samples and primers by modifying template configuration files. The specific functioning of the three workflows are described in more detail in the supplemental material and on the github page.

### PCR amplification and ASV generation

272 DNA samples from GEOTRACES cruises GA03 and GP13 were used to generate PCR amplicons, and represent either surface (GP13) or surface plus upper mesopelagic water samples (GA03). GP13 is a longitudinal transect in the southern Pacific, whereas GA03 is a longitudinal transect in the north Atlantic. DNA used for PCR amplicons was the exact same DNA material used to produce metagenomes described by Biller et al (2018) (20) except that it was diluted to 0.5 ng/µL in low-EDTA TE buffer prior to amplification. DNA was amplified with the 515Y/926R primers (515Y: 5’-GTGYCAGCMGCCGCGGTAA / 926R: 5’-CCGYCAATTYMTTTRAGTTT) (3) using 0.5ng of DNA template in a 25 µL reaction with a final primer concentration of 0.3mM. Primers were part of a larger construct with Illumina adapter constructs/barcodes already included, allowing for single-step library preparation. Thermocycling was as follows: an initial denaturation at 95°C for 120s followed by 30 amplification cycles of 95°C × 45s; 50°C × 45s; 68°C × 90s and a final elongation step at 68°C × 300s. Laboratory methods are described in more detail at https://dx.doi.org/10.17504/protocols.io.vb7e2rn. Both 16S and 18S rRNA mock communities (staggered and even versions of both (3, 13)) were included in the sequencing run as quality control.

Amplicons were sequenced with HiSeq 2x250 RapidRun technology at the USC sequencing center in a run pooled with shotgun metagenomic samples. Raw basecall data was demultiplexed with bcl2fastq (- r 20 -p 20 -w 32 --barcode-mismatches 0) using the 6 basepair reverse indices and subsequently with the forward 5 basepair barcode with cutadapt according to the templates provided at github.com/jcmcnch/demux-notes. Raw data from this step of the pipeline is available on NCBI under the umbrella Bioproject PRJNA659851 and three sub-projects PRJNA658608 (controls), PRJNA658384 (GA03), and PRJNA658385 (GP13).

Sequence data was analyzed with a custom *in-silico* pipeline available at github.com/jcmcnch/eASV-pipeline-for-515Y-926R and based on qiime2 2019.4 (45). Briefly, 16S and 18S rRNA sequences were first partitioned using a custom 16S/18S rRNA-specific databases derived from the SILVA 132 and PR2 (11, 46) which is available at osf.io/e65rs/. After partitioning, primers were removed from amplicons allowing up to 20% mismatches to the primer, and raw sequences were imported into qiime2 and denoised with DADA2 (10). Taxonomy was assigned using a naïve Bayesian classification algorithm with the SILVA 132 database subsetted to the amplicon region as a reference (47). Sequences identified as chloroplasts were re-annotated with PhytoRef (24), and 18S rRNA sequences were additionally annotated with PR2 (46). A step-by-step protocol for these informatic steps is available at https://dx.doi.org/10.17504/protocols.io.vi9e4h6. After denoising and annotation, biom tables were exported with taxonomy as tsv files and converted to relative abundances which were used as input for the metagenome-amplicon comparison.

## Acknowledgements

This work was supported by the Simons Collaboration on Computational Biogeochemical Modeling of Marine Ecosystems (CBIOMES) grant 549943 to JAF, the Gordon and Betty Moore Foundation Marine Microbiology Initiative grant 3779 and NSF grant OCE-1737409. We gratefully acknowledge the effort of the scientists who collected the BioGEOTRACES field samples analyzed here: Christel Hassler, Debbie Hulston, and Elizabeth Maas (GP13) and Jeremy Jacquot (GA03). We also acknowledge the efforts of Keven Dooley and Madeline Williams who conducted DNA extractions, which was funded in part by grants to Sallie W. Chisholm at MIT that enabled the collection and processing of DNA samples from GEOTRACES cruises. These include the Simons Foundation (Life Sciences Project Award ID 337262, S.W.C; SCOPE Award ID 329108, S.W.C), the Moore Foundation (Grant IDs GBMF495 and GBMF4511), and the National Science Foundation (OCE-1153588, OCE-1356460, and DBI-0424599). We thank S.W.C. for donating the samples and supporting parts of the study.

## Supplementary Information

### Supplementary Methods

#### Software Implementation

Software dependencies are handled by conda implemented in snakemake (1). Examples of how the pipeline can be run are available at github.com/jcmcnch/MGPrimerEval in the “runscripts” folder. Both the “classify” and “compare” workflows depend on the output of the “compute” workflow. The “classify” workflow is for detailed exploration of the taxonomy of retrieved sequences and the “compare” workflow allows for intercomparisons between SSU rRNA amplicon sequence variants (ASVs) and metagenomic SSU rRNA from the same amplicon region.

#### Compute workflow (“Snakefile-compute.smk”; Fig S30)

This portion of the analysis calculates the coverage of primer sequences for metagenomic samples specified in a configuration file. Results are calculated as the fraction of total SSU rRNA reads matching the primer at 0, 1, and 2-mismatches for 4 taxonomic partitions as described below. A rule graph summarizing the steps is shown in Figure S30. Briefly, fastq reads are first cleaned using fastp’s (v 0.14.1) default parameters and trimming 3 trailing and leading bases (2). To retrieve SSU rRNA sequences from metagenomes, we used the phyloFlash (v 3.3b1) package (3) which uses bbmap (v 38.69) (4) and a curated version of SILVA132 (5) to retrieve both 16S and 18S rRNA sequences. Since retrieved sequences contained a small fraction of low- complexity repeats that were not homologous to the SSU rRNA, we used komplexity (v 0.3.6) (6) to remove these likely spurious hits.

SSU rRNA sequences thus identified were then split into 4 different pools (Eukaryota, Archaea, Bacteria (non-cyanobacterial), and Cyanobacteria (including chloroplasts)) using bbsplit.sh v38.22 (4) comparing against curated subsets of the SILVA/PR2 reference databases (5, 7) (sequences and a template script for creating databases are available at osf.io/e65rs/). Partial SSU rRNA sequences were then aligned using pyNAST v1.2.2 (8) to group-specific references available as part of the github repository. Primer-specific subsets were then parsed from the alignment using python/biopython (9), only outputting sequences that overlap the primer region plus 5 leading or trailing bases. The 1-based coordinates of each primer region (for each organismal reference) must be supplied by the user; several widely-used primers are already included in template configuration files.

To place all reads in the correct orientation, fastq files were reverse-complemented based on information found in pyNAST alignment headers. cutadapt (version 1.18) (10) was used to identify primer regions in fastq sequences with 0, 1, 2, and 6-mismatches to the primer. Only the 0, 1 and 2- mismatch data were considered for plots/results following the established conventions from tools such as SILVA *testprime.* The 6-mismatch threshold was used to exclude any spurious matches caused by pyNAST misalignments or distantly homologous sequences; any sequence with greater than 6- mismatches to the primer was not further considered. Based on primer comparisons to references, this cutoff is likely to include even highly divergent taxa so long as they have a true primer binding site (NB: this is not true for some primers designed to specifically target a particular group - e.g. the 806RB primer does not contain a binding site for eukaryotic sequences). Primer coverage at each mismatch cutoff was then calculated using only sequences where all bases in the primer region plus 5 leading and trailing bases had a phred score of > 30. Plots shown in the main and supplemental text were generated with python/cartopy (11), ggplot2 (12), veusz (13), and snakemake (1) (Fig 1; Fig 2, Fig S1-S29; Fig 3; Fig S31-S32 respectively).

#### Classify workflow (“Snakefile-classify.smk”; Fig S31)

In order to determine which taxa were missed by a particular primer set and provide recommendations about how to further improve existing primers’ performance, we implemented an optional subsequent workflow to taxonomically classify reads and provide summary statistics for each group. Briefly, full- length fastq files from mismatched taxa (that is, SSU rRNA sequences with a primer-binding region but not matching primers at a given threshold) are classified with VSEARCH 2.10.4 (14) and the SINTAX algorithm (15) against the SILVA132 nr99 database (5) or PhytoRef (16) databases (for cyanobacteria/plastids) that had been converted into “.udb” format. The same procedure was followed for matches except a maximum of 5,000 reads were randomly subsampled using “filterbyname.sh” (4) to reduce computation time.

Summary statistics and output files were then generated for each unique combination of group, primer, and mismatch threshold (e.g. EUK, 926R, 0-mismatch) at order-level resolution since informative taxonomic annotations could generally not be made at finer taxonomic levels. Summaries contain the fraction of each taxon not matching the primer at a given mismatch threshold, as well as the fraction of total sequences represented by these mismatches. In addition, multiple sequence alignments of mismatches and their relative abundances are generated, with summary plots automatically generated showing the relative abundances of particular mismatches both for the whole dataset and subsetted by individual taxa. In addition, the same output files are generated for specific order-level taxa, allowing an investigator to see whether a given primer covers a particular taxon of interest. For the datasets analyzed in this study, raw output files/plots can be downloaded at https://osf.io/gr4nc/.

Our pipeline generates coverage information at the individual primer level since metagenomic reads will rarely/never cover the entire amplicon region (depending on lengths of sequence reads and the amplicon region). However, it is advantageous to present results in terms of predicted coverage for the primer set. If the mismatches to the forward and reverse primer were completely non-overlapping (i.e. the mismatches were to completely different organisms) then the mismatched fractions would be simply additive (“worst case” as mentioned in main text and shown in Fig 3). In contrast, if organisms overlapped between the forward and reverse primer, the largest mismatched fraction for that organism could be used. We used the taxonomic assignments discussed above to estimate this fraction of overlapping sequences (“rule normalize_summaries” and its dependent rules/scripts) and thus correct the primer coverage values as shown in Fig 2.

#### Compare workflow (“Snakefile-compare2tags-16s.smk” and “Snakefile-compare2tags-18s.smk”)

This workflow compares the abundance of taxa quantified by universal 16S and 18S rRNA amplicons using the 515Y/926R primers with abundances of taxa from metagenomic SSU rRNA in the same region of the molecule targeted by the PCR primers. The primer region is targeted by subsetting pyNAST alignments by primer coordinates. Only reads with at least 100bp overlap with the amplicon region are retained, and subsetted reads from the three prokaryotic categories are merged for the 16S rRNA comparison. For 18S rRNA reads that do not overlap (since the amplicon region is longer than the 2x250 sequencing length employed here), they were subsetted to the trim length of the partial amplicons (e.g. 220bp for forward read, 180 bp for reverse read).

These metagenomic sequences were then compared using BLASTn (17) (minimum 97% similarity, requiring 100% overlap) against a sample-specific database of exact amplicon sequence variants (ASVs) generated with DADA2 (18) that had a nonzero abundance in that particular sample. In order to make a direct comparison between the two data sources, the ASVs were clustered for each sample based on BLAST results with networkx (19) to account for the resolution mismatch between short metagenomic reads and longer, denoised amplicons. Total relative abundances of each cluster were plotted using seaborn (https://seaborn.pydata.org/) and sample-sample correspondence was evaluated using least squared linear regression with the stats.linregress function in scipy (20). The R^2^ value of this regression was used to evaluate general trends in MG-ASV sample-sample correspondence, and was plotted against both environmental covariates and the “worst-case” (simply additive) predicted primer performance (Fig 3b).

#### Amplicon Analysis Pipeline Supplementary Information

SSU rRNA partitioning was carried out with the database mentioned in the main text (osf.io/e65rs/) using bbsplit.sh (4). These databases were constructed from SILVA132 (5) and PR2 (7) by first clustering with CDHIT (21) at 95% identity to reduce total data volume and subsequently curated to remove sequences that were extremely long or short relative to the average sequence length.

Demultiplexing and trimming steps were carried out with cutadapt (10), and trimming prior to denoising (for 18S rRNA sequences) was carried out with bbduk.sh (4).

### Supplementary Results

#### Technical evaluation of pipeline

After retrieval, quality-filtering, and subsetting to the primer region, dozens to hundreds of rRNA sequences were recovered for each group, sample, read direction (fwd & rev reads), and primer combination (Table 1). Considering the pelagic ocean datasets introduced above and discussed in detail below (BiHoBa, TARA, MSPINA, MBARI, SPOT), we retrieved between 158-1148 primer-matching regions for *Bacteria* for each combination (medians within datasets; Table 1), whereas lower numbers were retrieved for the remaining three groups of *Archaea*, *Cyanobacteria*+Plastids and *Eukarya* (on average 17-109, 46-186 and 32-202 respectively) most likely due to their lower relative abundances in the environmental DNA. Among the primers, smaller number of sequences were identified for the two primers located near the end of the SSU rRNA molecule. For example, for Tara samples, a median of 455 sequences were retrieved per sample for 27F (close to end of molecule) for *Bacteria* vs. 1299 per sample for 338R (middle of molecule). Similarly, 58 eukaryotic sequences were retrieved per sample with 1510R vs. 136 per sample for 1389F.

With the exception of regions near the end of the SSU rRNA molecule, sequence coverage was mostly evenly distributed across the alignments to group-specific references. Some relatively small regions had lower coverage, most likely due to their absence in dominant environmental taxa. For example, the SAR11 isolate HTCC1062 is missing nucleotides (relative to the *E. coli* reference) corresponding to small portions of loops in the V1, V2, V3, and V9 regions (22). We note also that phyloFlash retrieved approximately double the total number of 18S rRNA sequences compared to a graftM (23) package based on an alignment of the greengenes database (data.ace.uq.edu.au/public/graftm/7/) which was employed in an earlier version of our pipeline. In addition, the coverage across the *S. cerevisae* 18S rRNA reference was much more even with phyloFlash-retrieved 18S rRNA vs. graftM. No such bias was noted for 16S rRNA sequences, suggesting this graftM package may be appropriate for 16S rRNA retrieval but not 18S rRNA.

#### Primer Limitations and Potential Improvements

Table S1 shows a summary of the top 3 most dominant mismatched orders for each primer and group for the BiHoBa dataset. These samples mostly represent surface and upper mesopelagic environments across all organismal size classes (Table 1), making it useful for exploring general patterns of primer performance for oceanic ecosystems. Among the most prominent mismatches to *Archaea* are those between *Nitrosopumilales* MGI *Archaea* and primers 341F or 1389F. Neither of these primers were specifically designed for capturing archaeal sequences but our results show they could be modified to include most *Archaea* as shown in Figs S1-S7 for 341F, and likely also for 1389F given that a universal primer with high coverage exists for the same general region (1392R). For other primers, the dominant mismatches include abundant groups such as MGII, and MGIII but we also note a smaller number of mismatches from the poorly-described *Aigarchaeales* and *Woesearchaeia*.

For *Bacteria*, many of the abundant mismatches were already discussed in the main text. Other patterns of particular note are mismatches to abundant members of the SAR86 clade and *Flavobacteriales* for 341F/338R primers. Together, these groups represent over 4% of the total sequences, meaning they would likely be desirable to correct in the future. Readers should note that the dominant taxa in Table S1 are the most abundant sequences, and there are a number of rare bacterial taxa that have very low coverage. These include *Poribacteria*, a number of members of the candidate phyla radiation or CPR, *Rubrobacterales* (*Actinobacteria*), and *Chlamydiales*. Interestingly, some members of these taxa (e.g. *Poribacteria*, CPR) had more than one mismatch to the 926R primer set, meaning they are likely very evolutionarily-divergent from other organisms and may require a more extensive primer redesign to quantify them fully. However, we note that these organisms are very rare in these datasets, so their absences are unlikely to bias overall community profiles in oxic marine environments and that the additional degeneracies proposed in the main text will increase coverage for some of these taxa.

With respect to the cyanobacterial/chloroplast fraction, while the 515Y/926R primer pair achieves nearly perfect coverage, the other primers generally have significant mismatches to one or more abundant chloroplast groups, including the *Mamelliophyceae*, *Prasinophyceae*, and *Prymnesiophyceae*. This means that care must be used in interpreting chloroplast results from some popular primer sets which may miss important taxa.

Among eukaryotic (nuclear 18S rRNA) sequences, the most commonly mismatched taxa included *Dinoflagellata*, *Discicristata, Protalveolata, Ochrophyta, Charophyta,* and *Metazoa.* As noted above, the universal primer sets appear to perform nearly as well or better versus *Eukarya-*specific primers.

Improvements described in the main text do increase 18S rRNA coverage for the 515Y/926R primers, but a fully comprehensive 18S rRNA primer may be less feasible since primer binding regions have a much higher sequence diversity versus 16S rRNA. This may be due to large copy numbers of SSU rRNA, some of which may become pseudogenized and undergo random evolutionary drift (24). In practical terms, this means that some of these more rare variants will inevitably be missed by primers since creating an overly-degenerate primer could cause problems with non-specific amplification during PCR. The same may be true for some mitochondrial 16S rRNA genes that have become pseudogenized after being incorporated into the host nuclear genome (25).

#### A Practical Consideration for the use of the 515Y/926R primer set

One concern about using truly universal primers is the amplification of non-microbial sequences, e.g. in the case of host-associated microbiomes. This has led to the design of prokaryotic-specific primer sets (e.g. 515Y/806RB), that intentionally do not amplify 18S rRNA sequences. However, universal primers retain the advantage of quantifying a broad swathe of the entire biological community together. Therefore, such primers could be potentially be very useful if 18S rRNA sequences do not overwhelm the SSU rRNA sequences in typical samples.

To test this, we quantified the total fraction of 18S rRNA (out of the total 16S + 18S rRNA) for the unfractionated (>0.2µM) BiHoBa metagenomes. For these data, we show 18S rRNA sequences rarely exceed ∼1/3 of the total SSU rRNA sequences (Fig S29), indicating that universal primers will still be useful even when whole seawater samples are analyzed, providing quantification of both prokaryotic and eukaryotic communities simultaneously. The fraction of eukaryotic SSU rRNA relative to total 16S + 18S rRNA averaged 19.5 +/- 0.09 % (range = 0.024 - 0.7). Samples with large fractions of eukaryotes (>0.25) were dominated by *Metazoa*, mainly composed of taxa such as *Arthropoda, Platyhelminthes,* and *Hydrozoa.* These animal-derived sequences have the potential to overwhelm the signal from other environmental eukaryotes, especially for unfractionated samples. This can be alleviated to some extent by increasing overall sequence coverage such that sufficient non-metazoan 18S rRNA sequences can also be recovered.

---

**Table S1:**
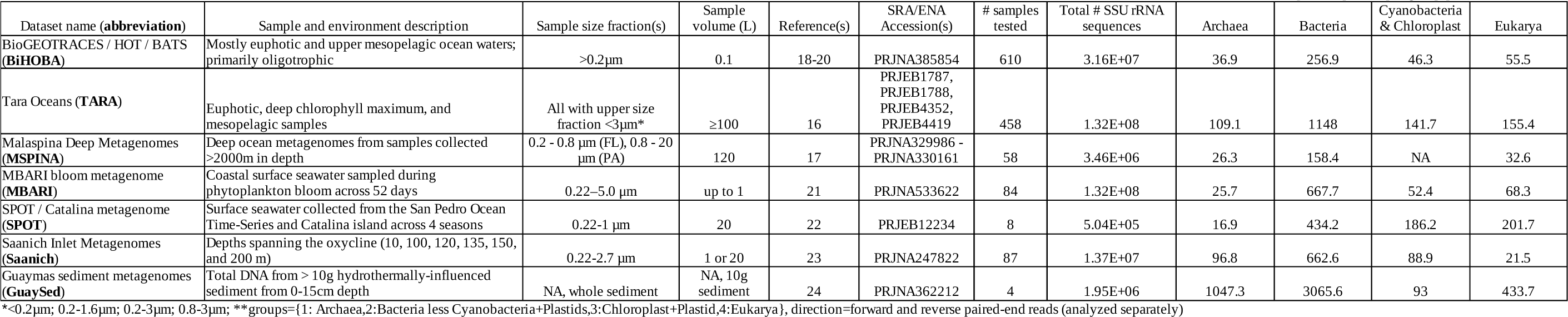
Data used in this study. Exact SRA/ENA Accessions available on github.com/jcmcnch/MGPrimerEval in config files.

| Dataset name (**abbreviation**) | Sample and environment description | Sample size fraction(s) | Sample volume (L) | Reference(s) | SRA/ENA Accession(s) | # samples tested | Total # SSU rRNA sequences | Median filtered sequences per sample/group/primer/direction** |  |  |  |
| --- | --- | --- | --- | --- | --- | --- | --- | --- | --- | --- | --- |
|  |  |  |  |  |  |  |  | Archaea | Bacteria | Cyanobacteria & Chloroplast | Eukarya |
| BioGEO TRACES / HOT / BATS (**BiHOBA**) | Mostly euphotic and upper mesopelagic ocean waters; primarily oligotrophic | >0.2 µm | 0.1 | 18-20 | PRJNA385854 | 610 | 3.16E+07 | 36.9 | 256.9 | 46.3 | 55.5 |
| Tara Oceans (**TARA**) | Euphotic, deep chlorophyll maximum, and mesopelagic samples | All with upper size fraction <3 µm* | ≥100 | 16 | PRJEB1787, PRJEB1788, PRJEB4352, PRJEB4419 | 458 | 1.32E+08 | 109.1 | 1148 | 141.7 | 155.4 |
| Malaspina Deep Metagenomes (**MSPINA**) | Deep ocean metagenomes from samples collected >2000m in depth | 0.2 - 0.8 µm (FL), 0.8 - 20 µm (PA) | 120 | 17 | PRJNA329986 - PRJNA330161 | 58 | 3.46E+06 | 26.3 | 158.4 | NA | 32.6 |
| MBARI bloom metagenome (**MBARI**) | Coastal surface seawater sampled during phytoplankton bloom across 52 days | 0.22–5.0 µm | up to 1 | 21 | PRJNA533622 | 84 | 1.32E+08 | 25.7 | 667.7 | 52.4 | 68.3 |
| SPOT / Catalina metagenome (**SPOT**) | Surface seawater collected from the San Pedro Ocean Time-Series and Catalina island across 4 seasons | 0.22-1 µm | 20 | 22 | PRJEB12234 | 8 | 5.04E+05 | 16.9 | 434.2 | 186.2 | 201.7 |
| Saanich Inlet Metagenomes (**Saanich**) | Depths spanning the oxycline (10, 100, 120, 135, 150, and 200 m) | 0.22-2.7 µm | 1 or 20 | 23 | PRJNA247822 | 87 | 1.37E+07 | 96.8 | 662.6 | 88.9 | 21.5 |
| Guaymas sediment metagenomes (**GuaySed**) | Total DNA from > 10g hydrothermally-influenced sediment from 0-15cm depth | NA, whole sediment | NA, 10g sediment | 24 | PRJNA362212 | 4 | 1.95E+06 | 1047.3 | 3065.6 | 93 | 433.7 |

\*<0.2µm; 0.2-1.6µm; 0.2-3µm; 0.8-3µm; \*\*groups={1: Archaea,2:Bacteria less Cyanobacteria+Plastids,3:Chloroplast+Plastid,4:Eukarya}, direction=forward and reverse paired-end reads (analyzed separately)

**Table S2:**
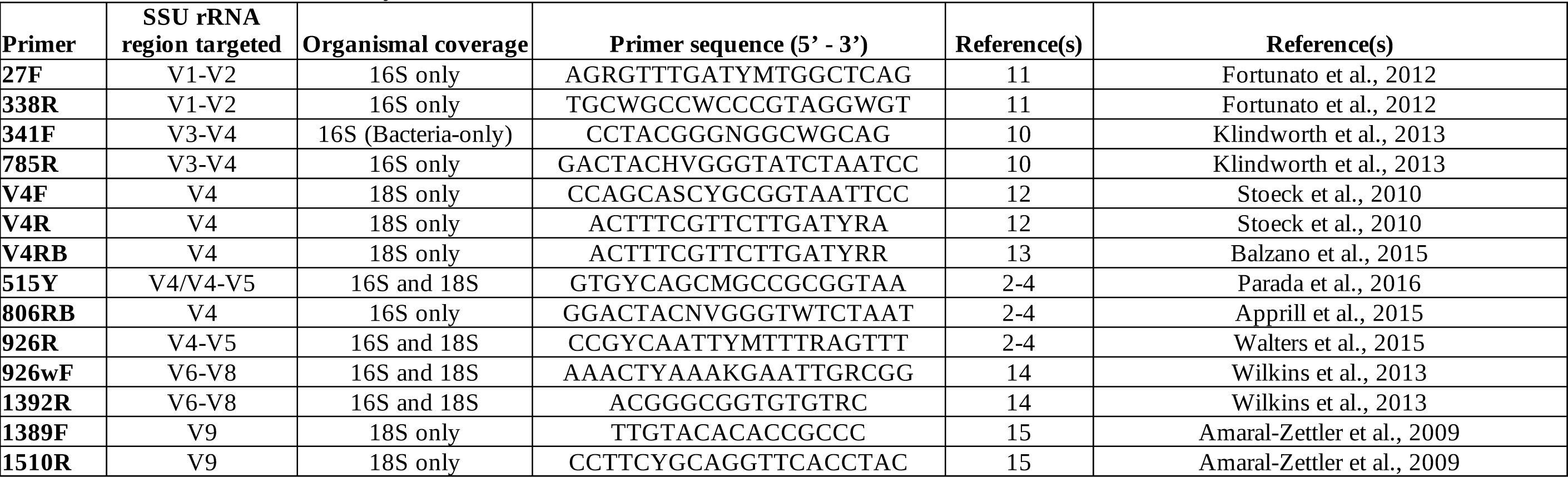
Primers tested in this study

**Table S3:** Top 3 most abundant mismatched orders for each organismal category in the BioHOBA dataset. Values in parentheses represent the fraction of sequences that do not perfectly match the primer at a 0- mismatch threshold as a fraction of total SSU rRNA sequences (matched + mismatched). For example, for the primer 785R, mismatches to SAR11 primer-binding regions represent 40% of total identified bacterial sequences (excluding cyanobacteria + plastids), whereas the most abundant mismatched taxon for 515Y is Rickettsiales at 0.2% total bacterial SSU rRNA read abundance. Primers that have “NA” for particular organismal category did not have a recognizable primer-binding region. This often represents an intentional design choice to avoid what was considered off-target amplification. For example, the 806RB primer was designed not to amplify 18S (Eukaryote) sequences.

| Primer | Archaea | Bacteria (without Cyanobacteria and plastid 16S) | Cyanobacteria (including plastid 16S) | Eukarya (18S nuclear SSU rRNA only) |
| --- | --- | --- | --- | --- |
| 27F | NA, no apparent primer binding site | Rickettsiales (0.008), Ectothiorhodospirales (0.008), SAR11 clade (0.002) | Prasinococcales (0.011), Cyanobacteria;Subsection I (0.003), Chrysophyceae-Synurophyceae (0.002) | NA, no apparent primer binding site |
| 338R | NA, no apparent primer binding site | SAR86 clade (0.027), Flavobacteriales (0.015), SAR11 clade (0.009) | Prymnesiophyceae (0.123), Bacillariophyta (0.006), Prasinococcales (0.005) | NA, no apparent primer binding site |
| 341F | Nitrosopumilales (0.51), MG II (0.41), MG III (0.07) | SAR86 clade (0.028), Flavobacteriales (0.016), SAR11 clade (0.016) | Prymnesiophyceae (0.121), Bacillariophyta (0.006), Prasinococcales (0.005) | NA, 5-mismatch to reference |
| 785R | MG II (0.003), Nitrosopumilales (0.002), MG III (0.0004) | SAR11 clade (0.40), Marinimicrobia (SAR406_clade); unclassified order/class (0.006), Verrucomicrobiae;Arctic97B-4 (0.003) | Mamiellophyceae (0.103), Prymnesiophyceae (0.006), Cyanobacteria;SubsectionI (0.001) | NA, 3-mismatch to reference |
| V4F | NA, 5-mismatch to reference | NA, 3-mismatch to reference | NA, 4-mismatch to reference | Dinoflagellata (0.022), Metazoa (Animalia) (0.010), Protalveolata (0.007) |
| V4R | NA, 8-mismatch to reference | NA, 7-mismatch to reference | NA, many mismatches to reference | Dinoflagellata (0.081), Discicristata (0.027), Protalveolata (0.026) |
| V4RB | NA, 8-mismatch to reference | NA, 7-mismatch to reference | NA, many mismatches to reference | Dinoflagellata, (0.040), Discicristata (0.013), Protalveolata (0.012) |
| 515Y | Nitrosopumilales (0.003), MG II (0.002), MG III (0.0003) | Rickettsiales (0.002), SAR11 clade (0.001), Chlamydiales (0.0006) | Cyanobacteria;Subsection I (0.003), Prymnesiophyceae (0.001), Pelagophyceae (0.000) | Dinoflagellata (0.021), Discicristata (0.014), Metazoa (Animalia) (0.010) |
| 806RB | Nitrosopumilales (0.002), MG II (0.002), Aigarchaeales (0.001) | Marinimicrobia (SAR406_clade); unclassified order/class (0.006), Flavobacteriales (0.005), SAR202 clade (0.002) | Mamiellophyceae (0.108), Prymnesiophyceae (0.014), Prasinophyceae (0.013) | NA, many mismatch to reference |
| 926R | Nitrosopumilales (0.016), MG II (0.001), Woeseearchaeia; unclassified order (0.0007) | Ectothiorhodospirales (0.007), Rickettsiales (0.004), SAR11 clade (0.002) | Cyanobacteria;Subsection I (0.003), Prymnesiophyceae (0.000), Dictyochophyceae (0.000) | Dinoflagellata (0.015), Protalveolata (0.012), Metazoa (Animalia) (0.010) |
| 926wF | Nitrosopumilales (0.016), MG II (0.001), Woeseearchaeia; unclassified order (0.0007) | Rhodospirillales (0.041), Ectothiorhodospirales (0.007), Rickettsiales (0.004) | Cyanobacteria;Subsection I (0.003), Prymnesiophyceae (0.000), Dictyochophyceae (0.000) | Metazoa (Animalia) (0.040), Ochrophyta (0.033), Dinoflagellata (0.016) |
| 1392R | Nitrosopumilales (0.002), Woeseearchaeia; unclassified order (0.002), MG II (0.001) | SAR11 clade (0.003), Clostridiales (0.001), Rickettsiales (0.001) | Mamiellophyceae (0.227), Dictyochophyceae (0.011), Bacillariophyta (0.002) | Dinoflagellata (0.014), Protalveolata (0.010), Metazoa (Animalia) (0.008) |
| 1389F | Nitrosopumilales (0.53), MG II (0.37), MG III (0.05) | Dadabacteriales (0.002), Clostridiales (0.002), SAR11 clade (0.002) | Mamiellophyceae (0.167), Cyanobacteria;Subsection I (0.103), Trebouxiophyceae (0.055) | Dinoflagellata (0.013), Protalveolata (0.008), Metazoa (Animalia) (0.006) |
| 1510R | NA, 4-mismatch to reference | NA, 4-mismatch to reference | Prasinophyceae (0.393), Pelagophyceae (0.375), Pavlovophyceae (0.054) | Metazoa (Animalia) (0.055), Charophyta (0.036), Protalveolata (0.012) |

**Table S4:**
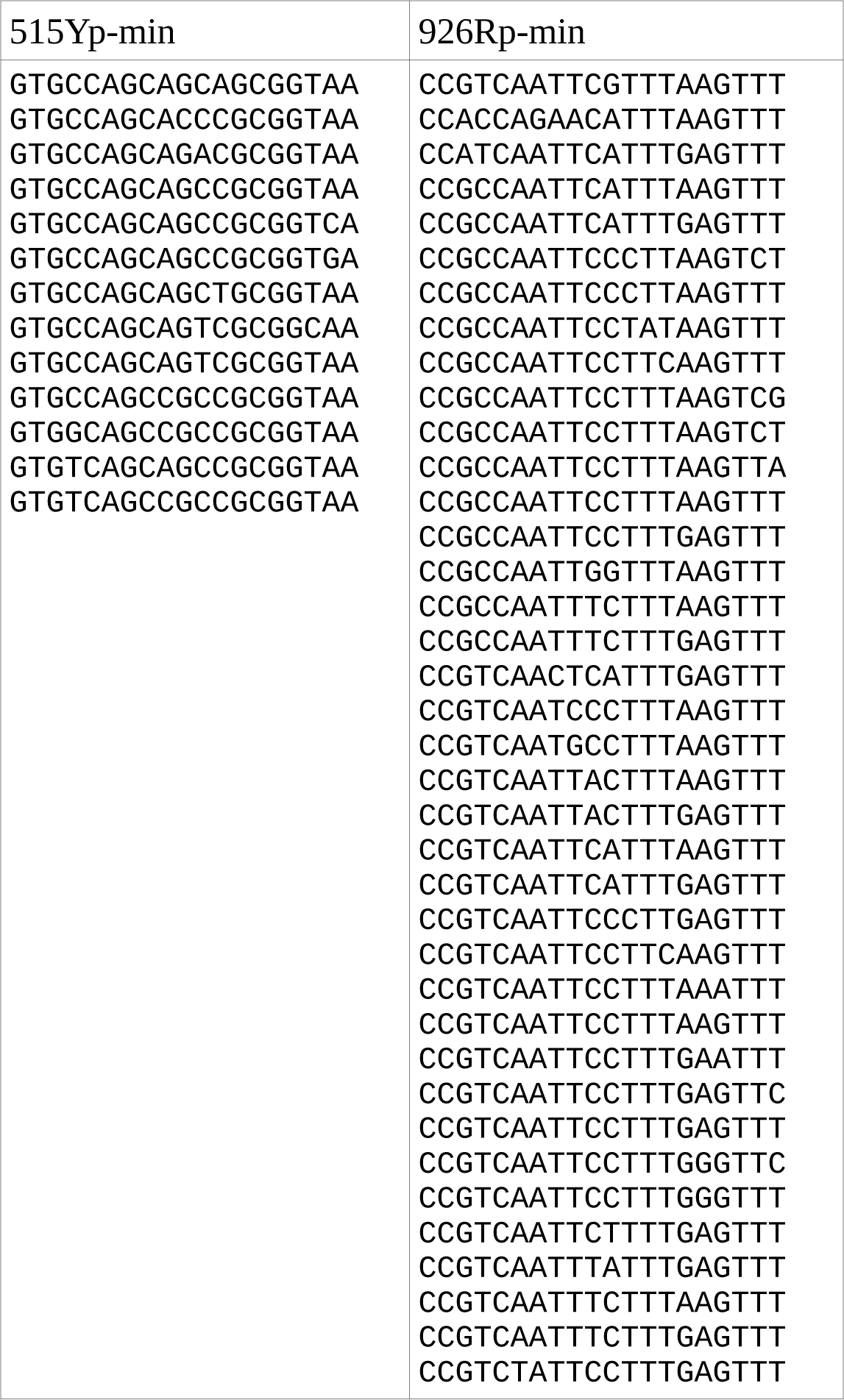
Oligonucleotide mixtures specified for the new primers 515Yp-min and 926Rp-min.

**Fig S1:**
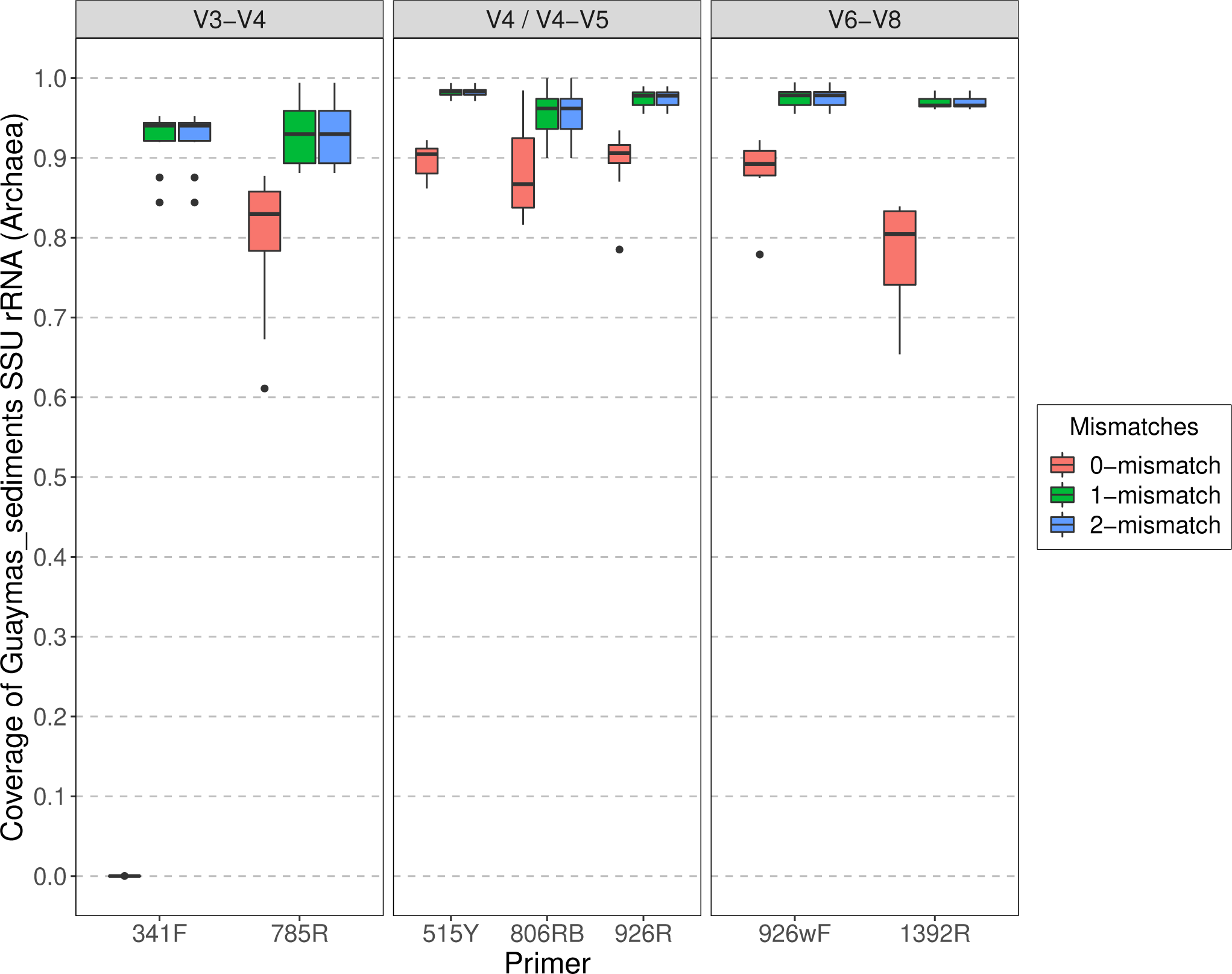
Predicted individual primer coverage allowing 0, 1, and 2-mismatches for *Archaea* using metagenomes derived from Guaymas Basin sediments.

**Fig S2:**
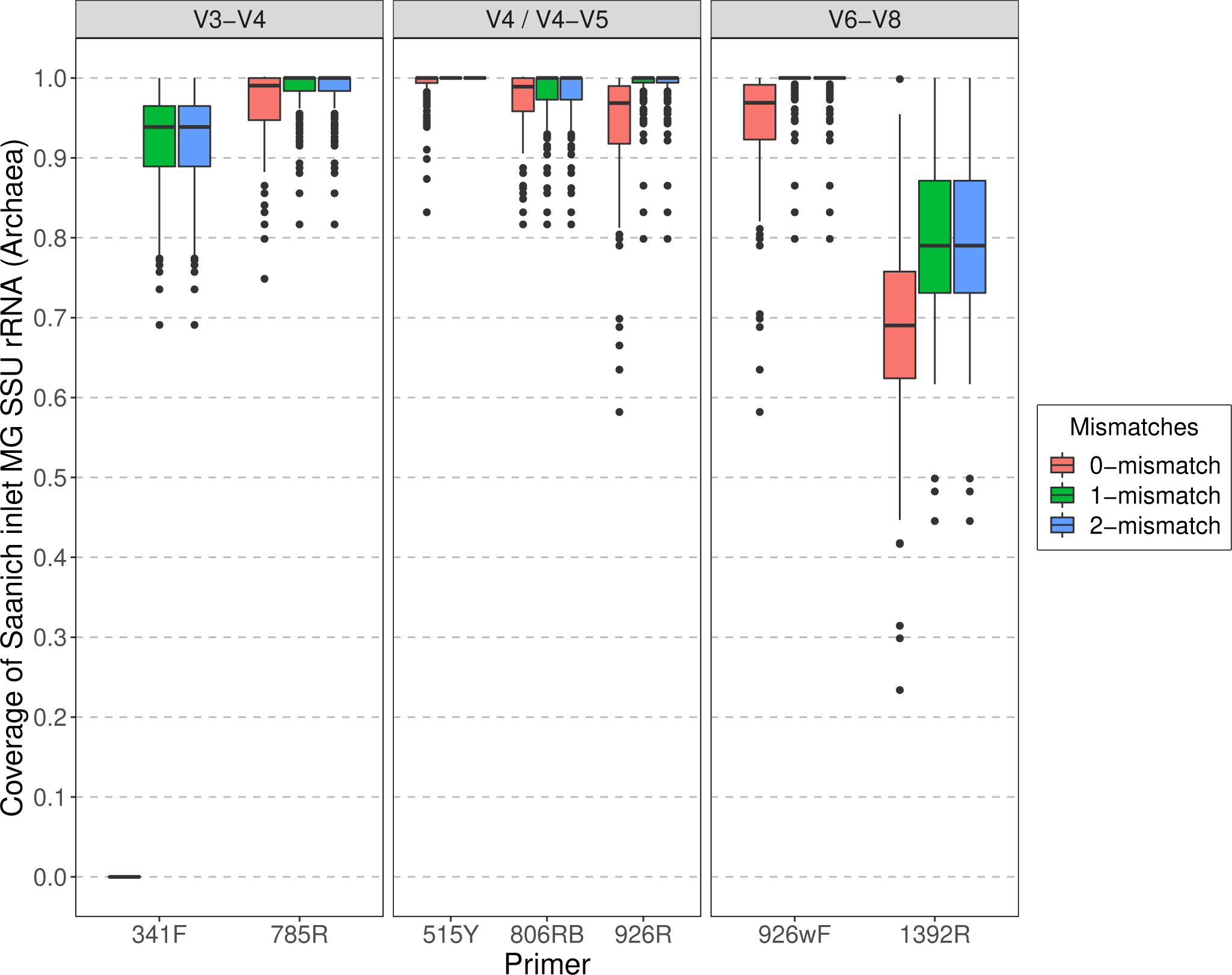
Predicted individual primer coverage allowing 0, 1, and 2-mismatches for *Archaea* using metagenomes derived from Saanich Inlet water samples.

**Fig S3:**
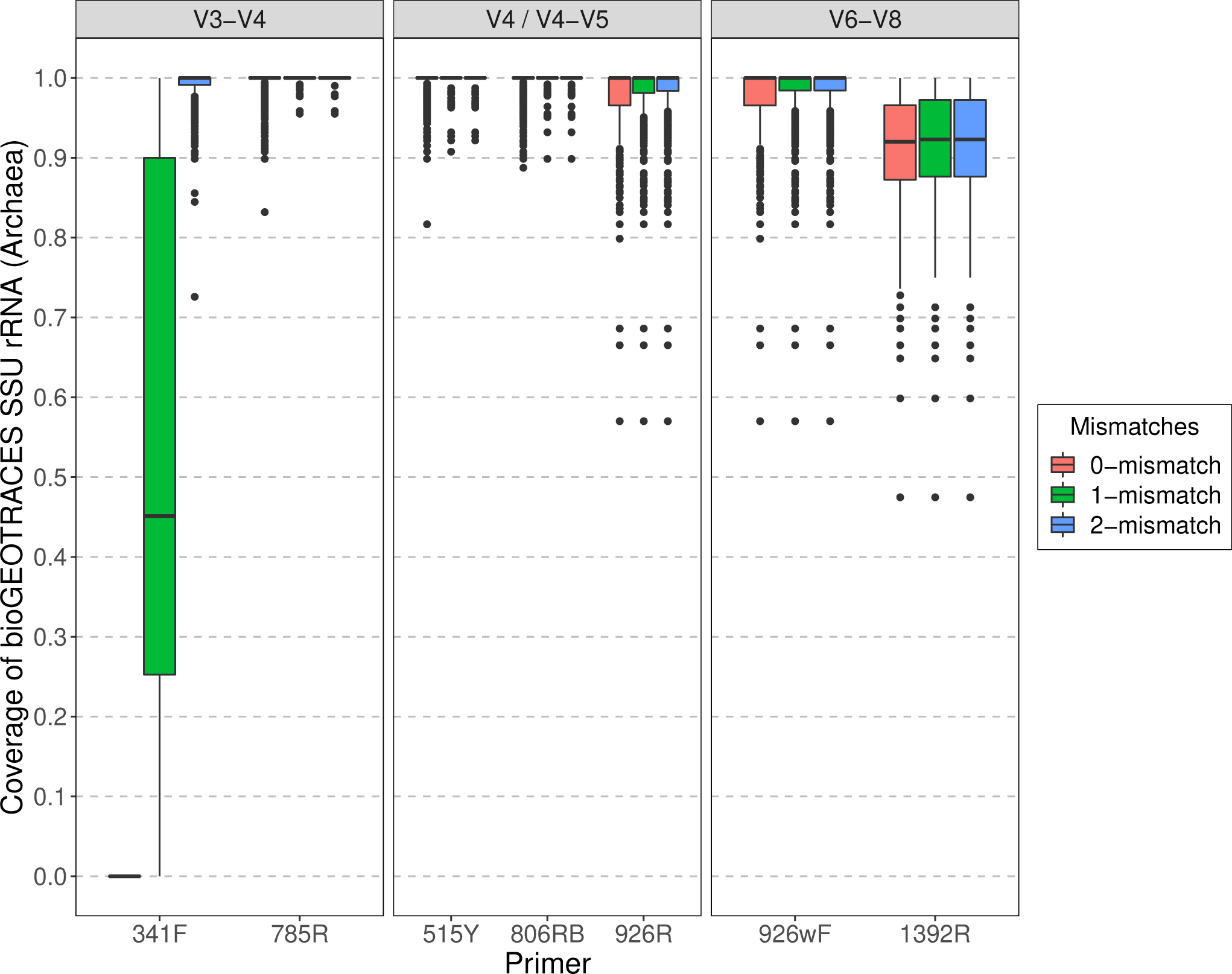
Predicted individual primer coverage allowing 0, 1, and 2-mismatches for *Archaea* using metagenomes derived from BioGEOTRACES/BATS/HOT water samples.

**Fig S4:**
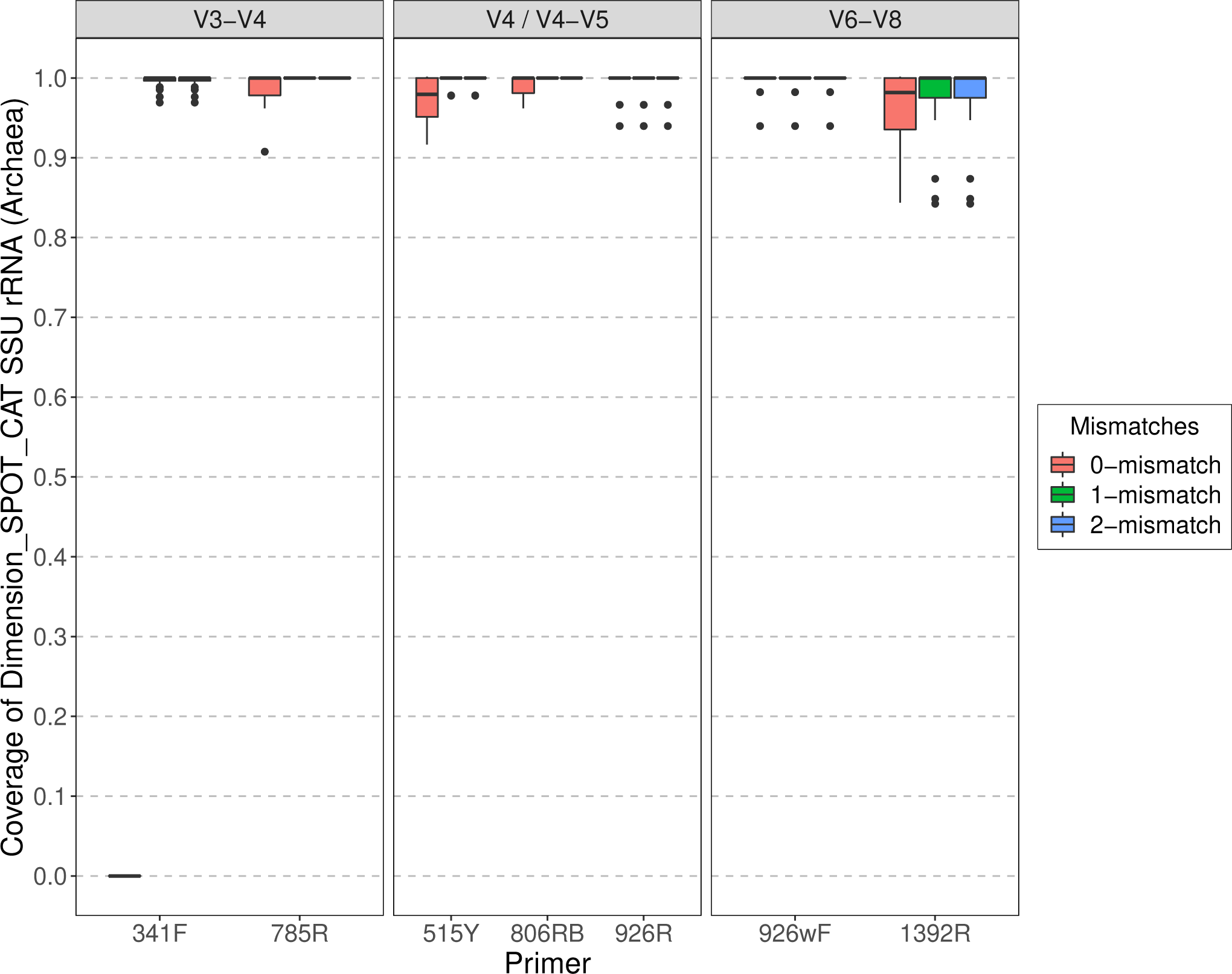
Predicted individual primer coverage allowing 0, 1, and 2-mismatches for *Archaea* using metagenomes derived from SPOT/Catalina water samples.

**Fig S5:**
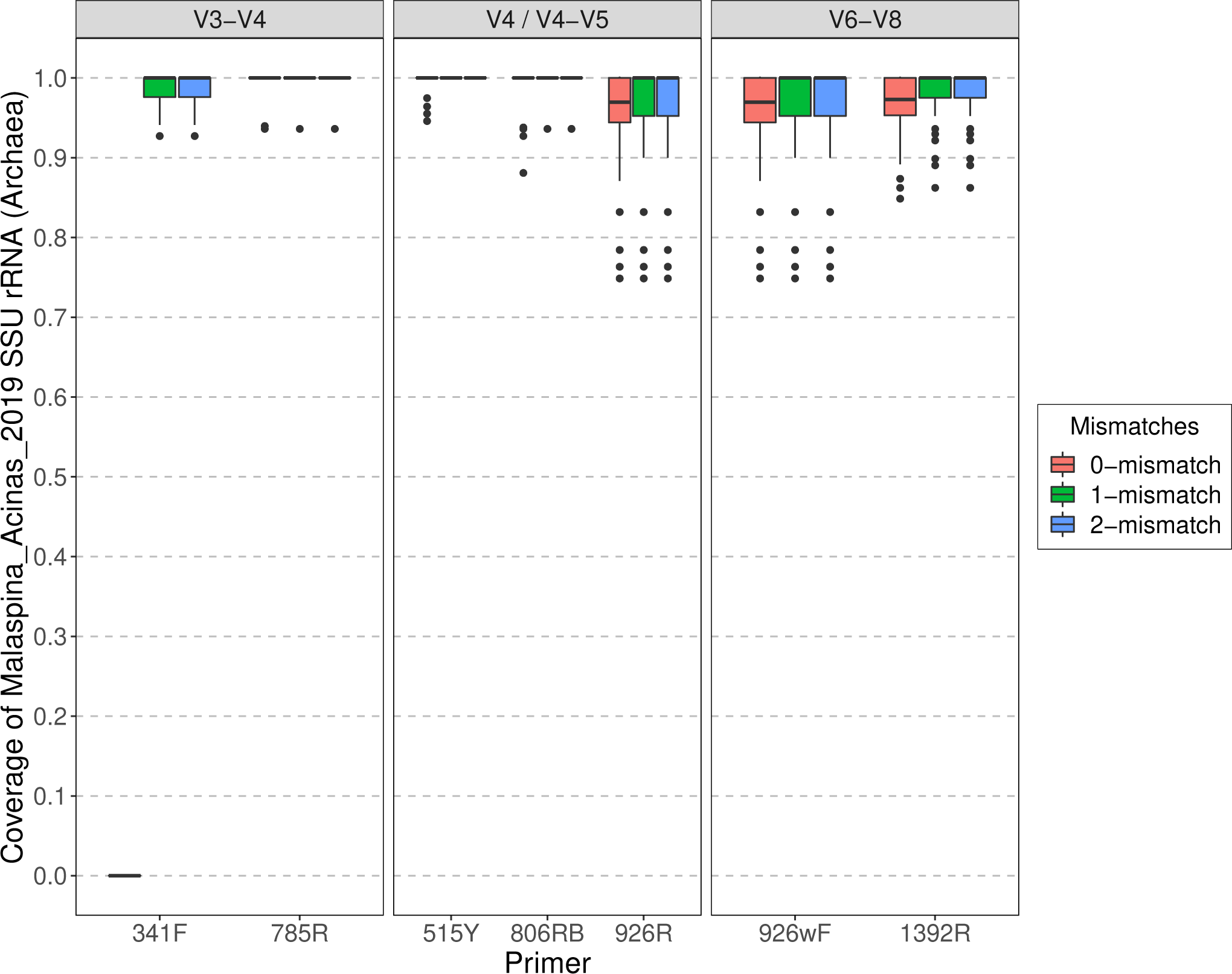
Predicted individual primer coverage allowing 0, 1, and 2-mismatches for *Archaea* using metagenomes derived from Malaspina deep-sea water samples.

**Fig S6:**
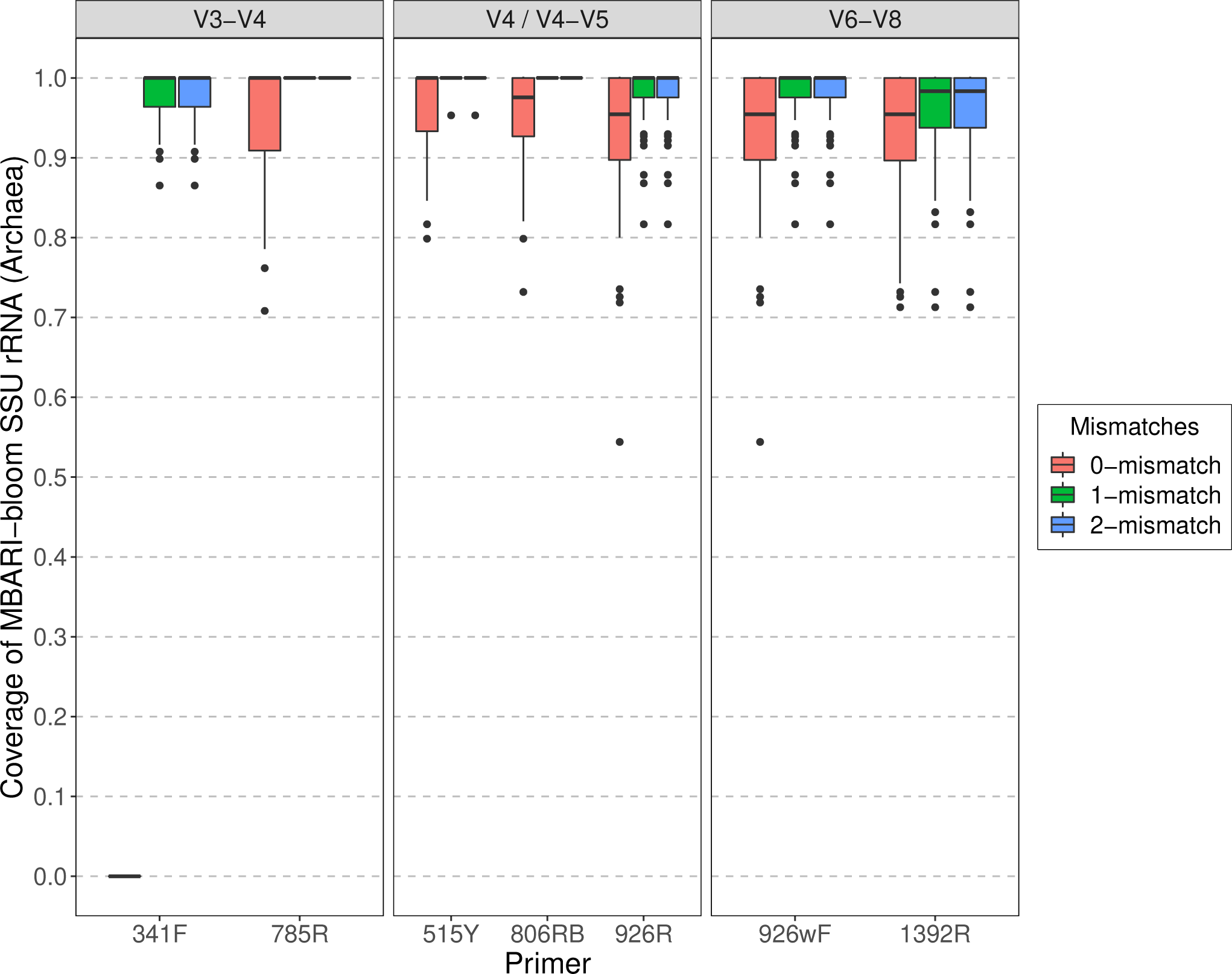
Predicted individual primer coverage allowing 0, 1, and 2-mismatches for *Archaea* using metagenomes derived from MBARI bloom water samples.

**Fig S7:**
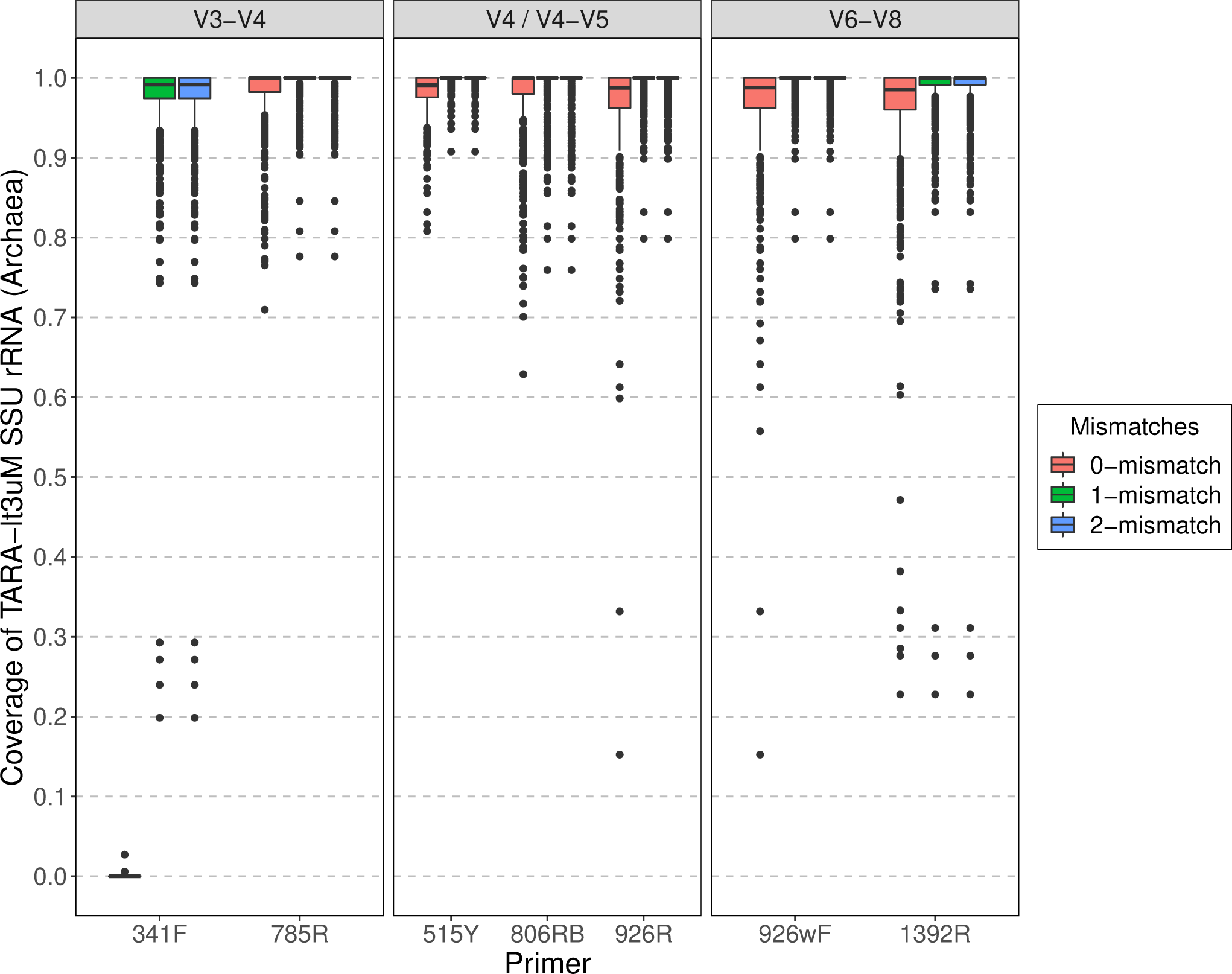
Predicted individual primer coverage allowing 0, 1, and 2-mismatches for *Archaea* using metagenomes derived from Tara Oceans water samples (<3µm).

**Fig S8:**
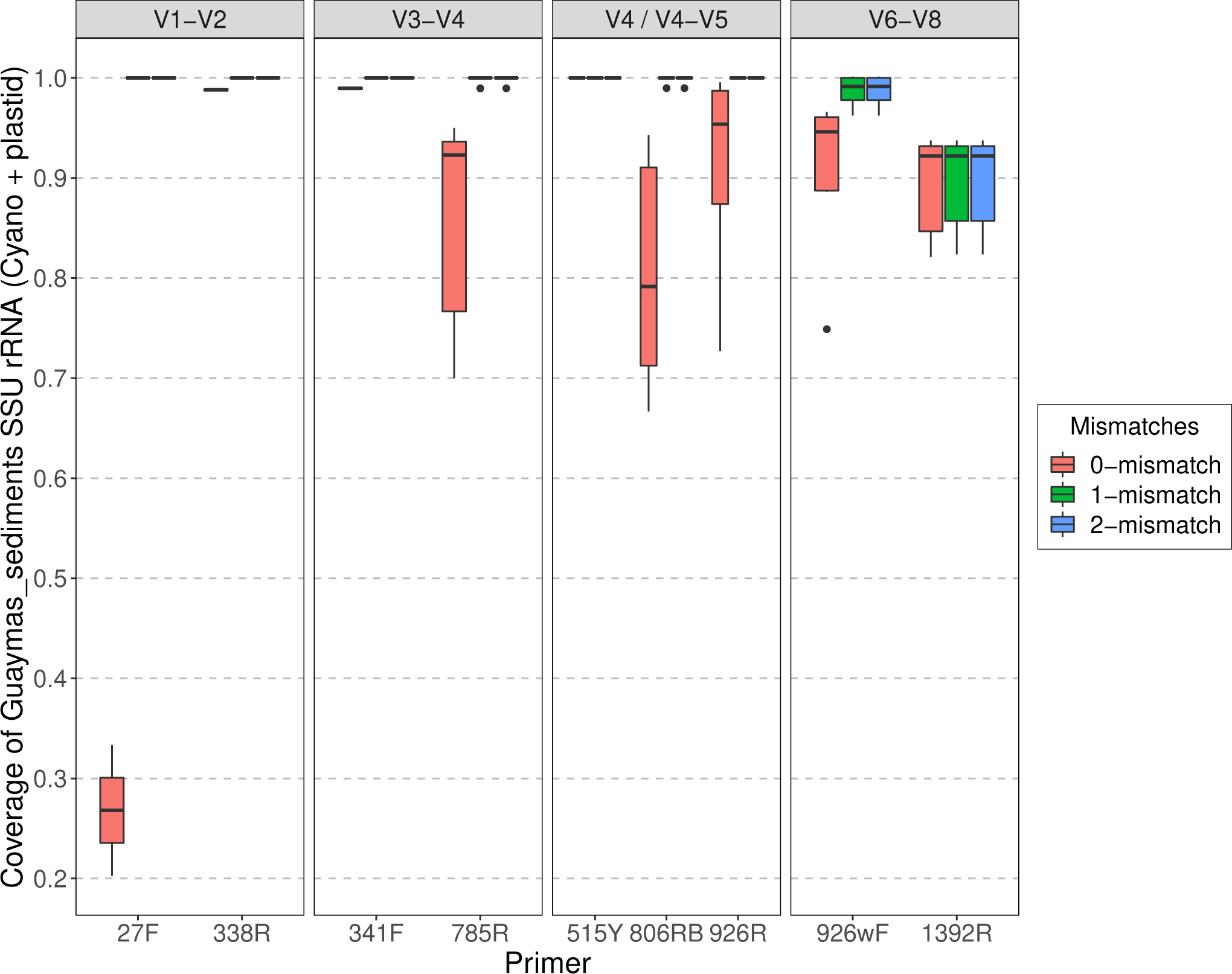
Predicted individual primer coverage allowing 0, 1, and 2-mismatches for *Cyanobacteria* and plastids using metagenomes derived from Guaymas Basin sediments.

**Fig S9:**
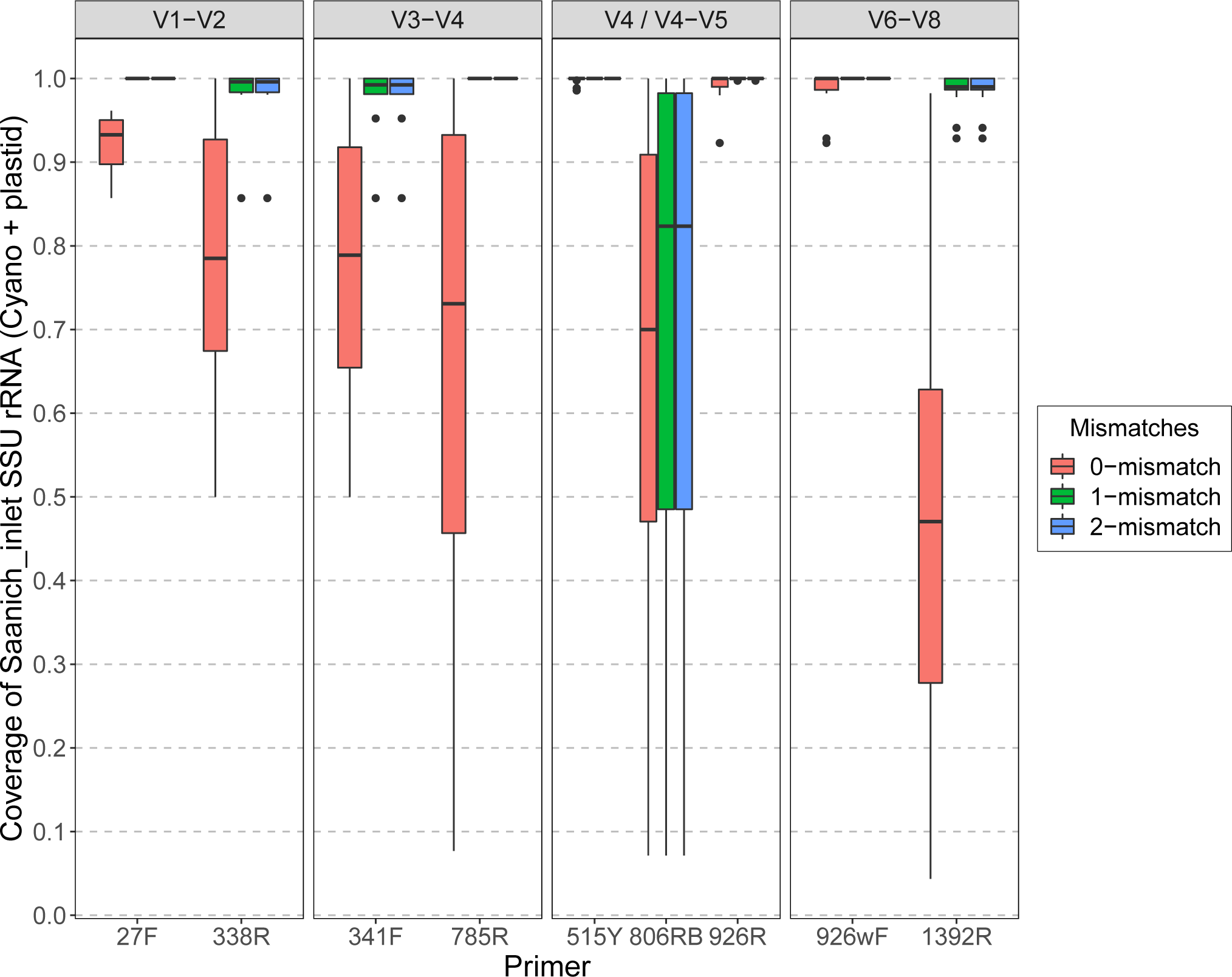
Predicted individual primer coverage allowing 0, 1, and 2-mismatches for *Cyanobacteria* and plastids using metagenomes derived from Saanich Inlet water samples.

**Fig S10:**
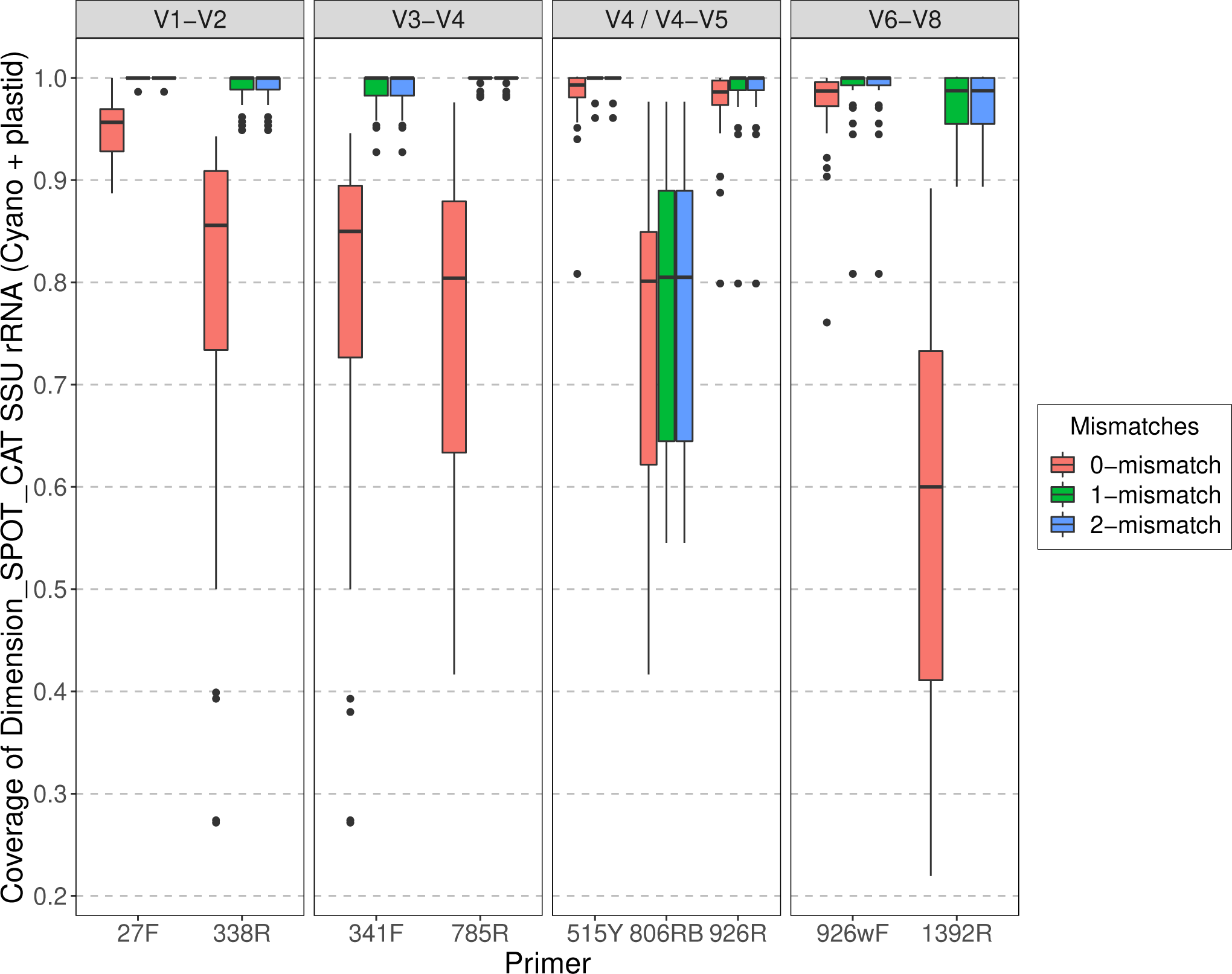
Predicted individual primer coverage allowing 0, 1, and 2-mismatches for *Cyanobacteria* and plastids using metagenomes derived from SPOT/Catalina water samples.

**Fig S11:**
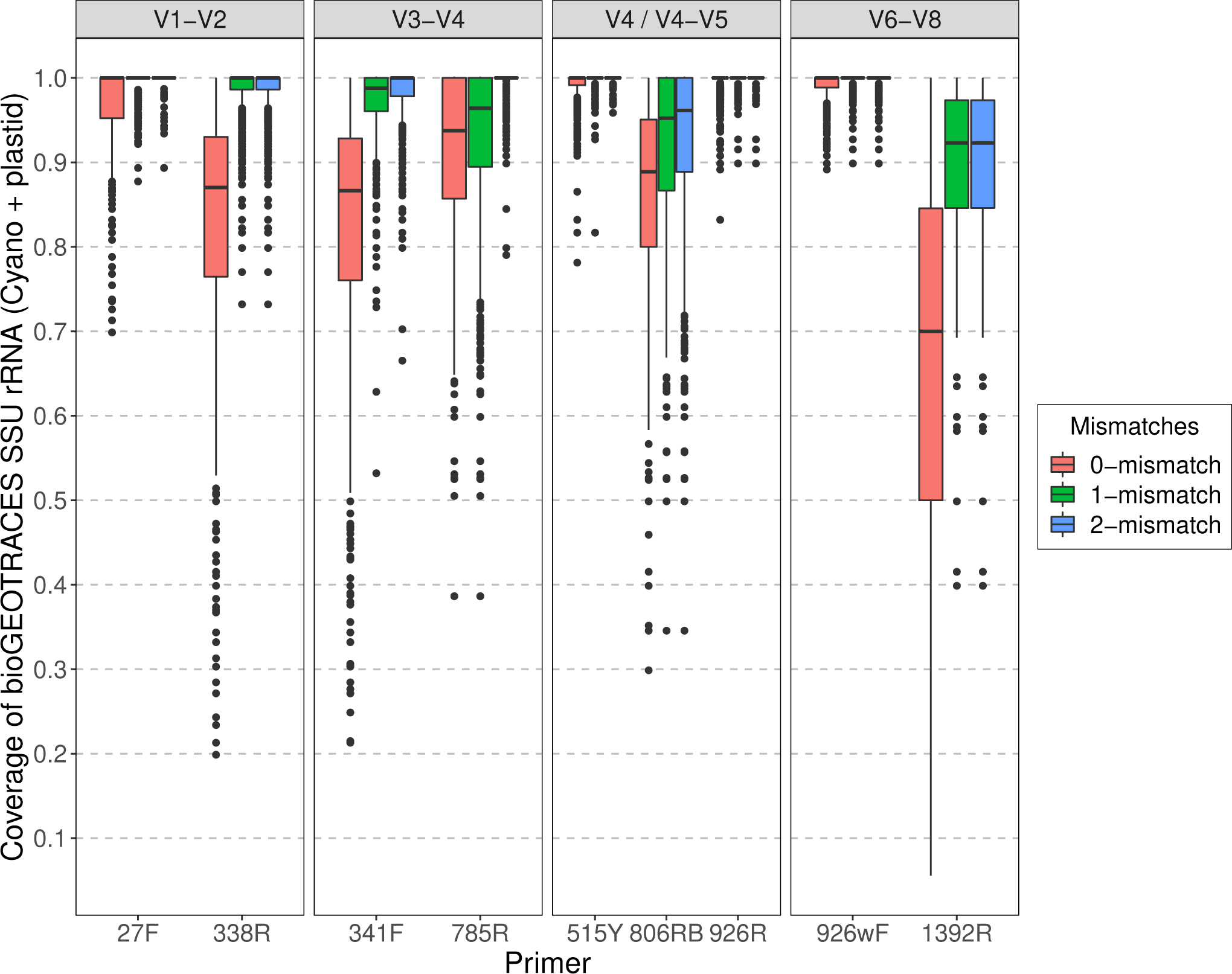
Predicted individual primer coverage allowing 0, 1, and 2-mismatches for *Cyanobacteria* and plastids using metagenomes derived from BioGEOTRACES/BATS/HOT water samples.

**Fig S12:**
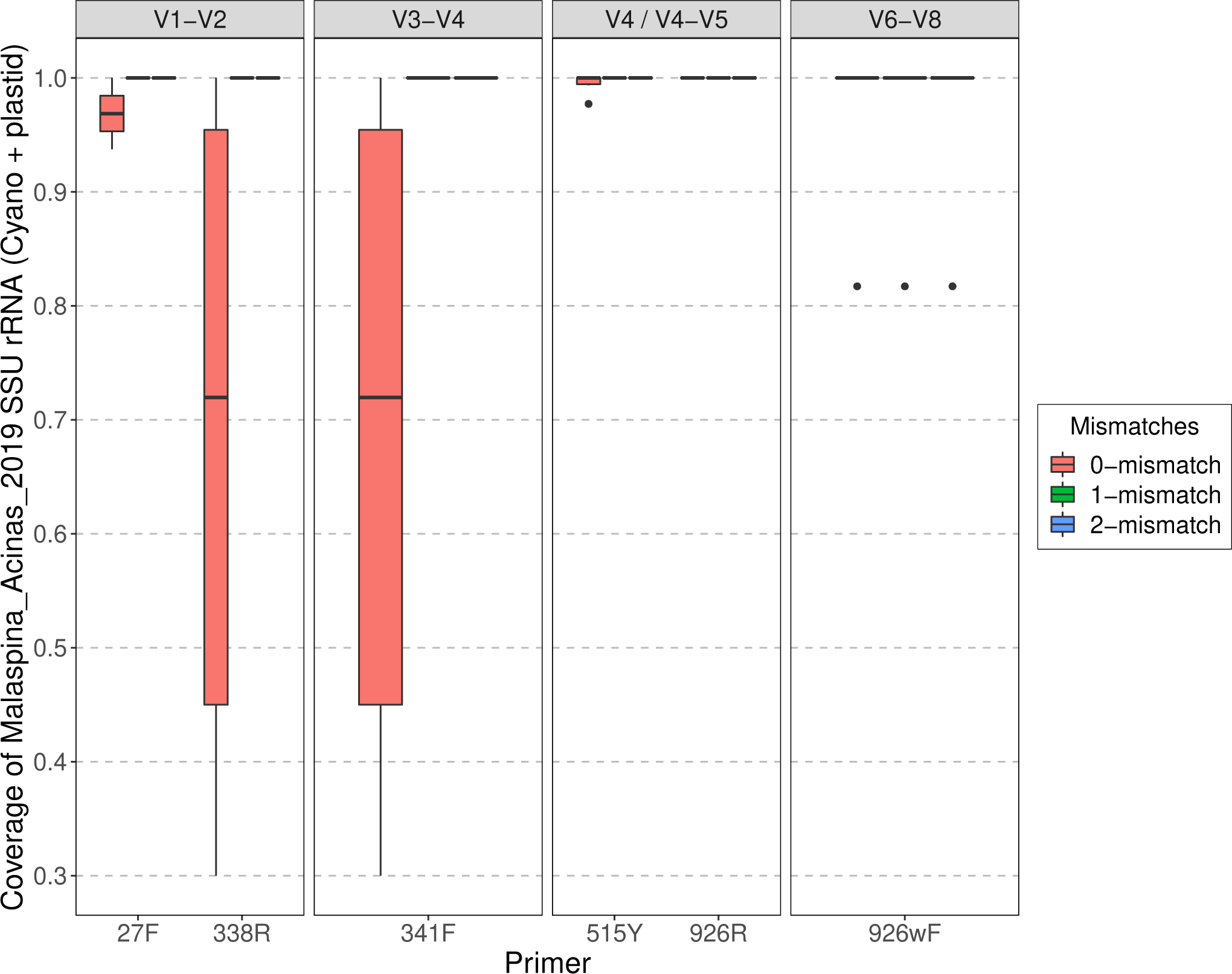
Predicted individual primer coverage allowing 0, 1, and 2-mismatches for *Cyanobacteria* and plastids using metagenomes derived from Malaspina deep-sea water samples.

**Fig S13:**
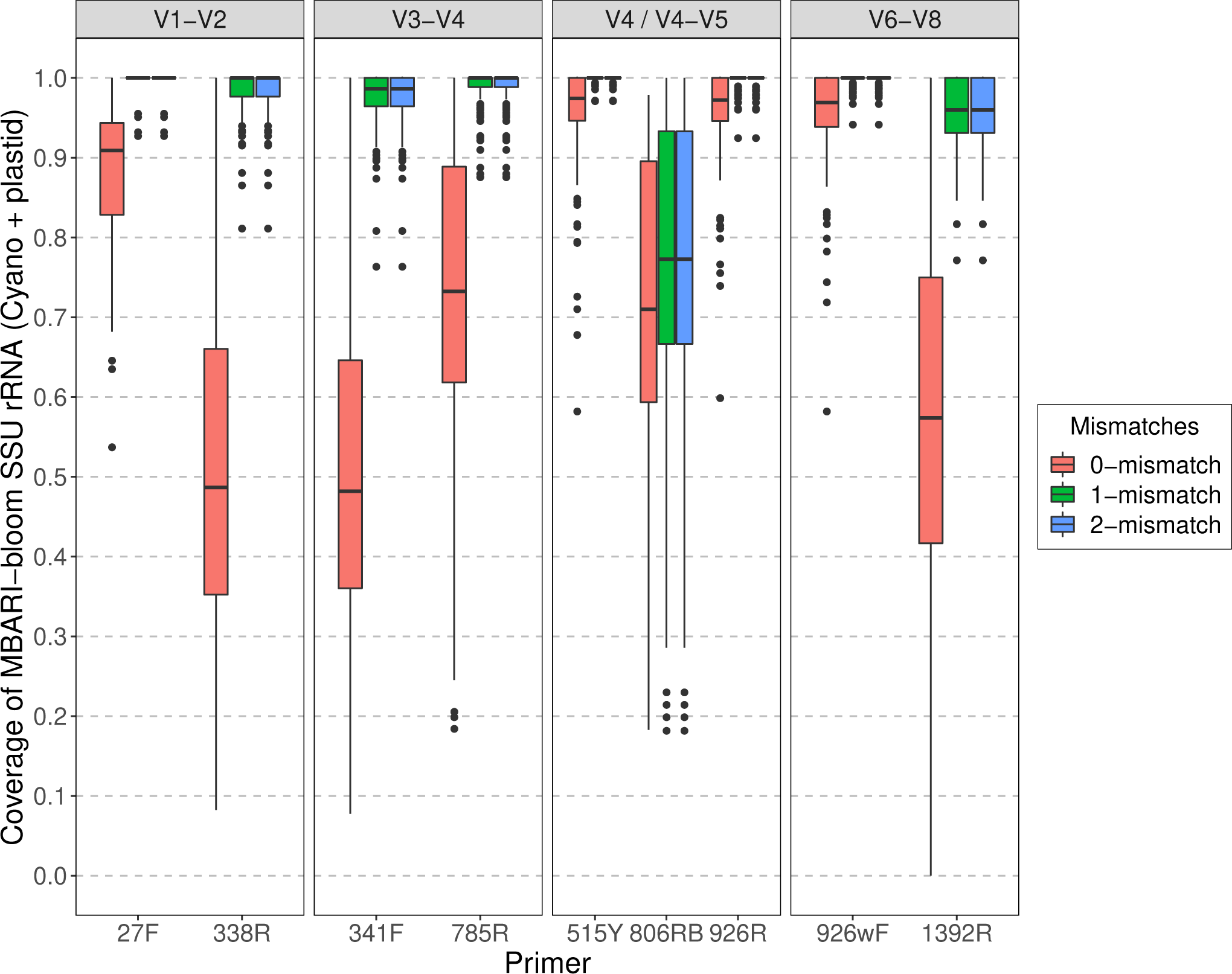
Predicted individual primer coverage allowing 0, 1, and 2-mismatches for *Cyanobacteria* and plastids using metagenomes derived from MBARI bloom water samples.

**Fig S14:**
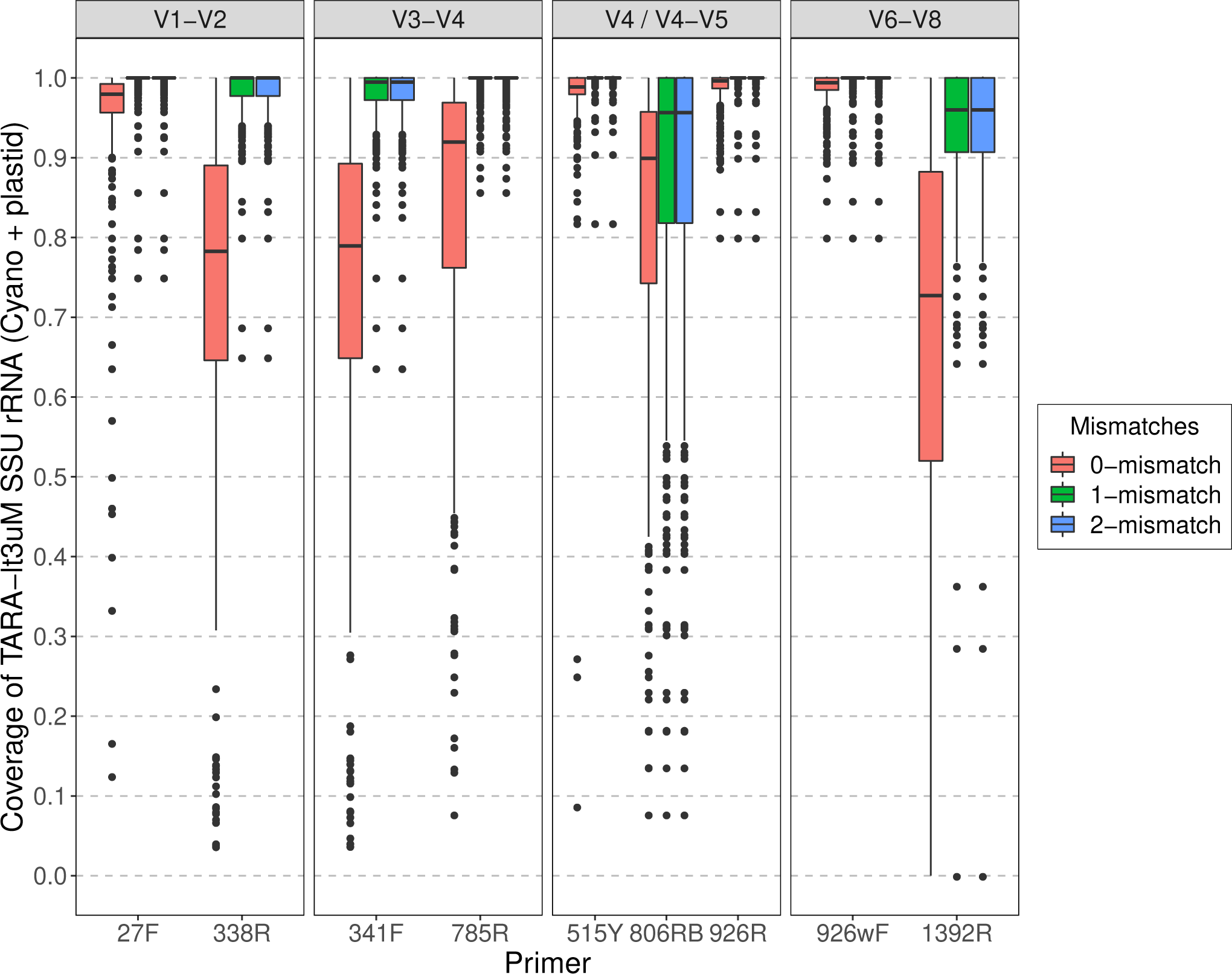
Predicted individual primer coverage allowing 0, 1, and 2-mismatches for *Cyanobacteria* and plastids using metagenomes derived from Tara Oceans water samples (<3µm).

**Fig S15:**
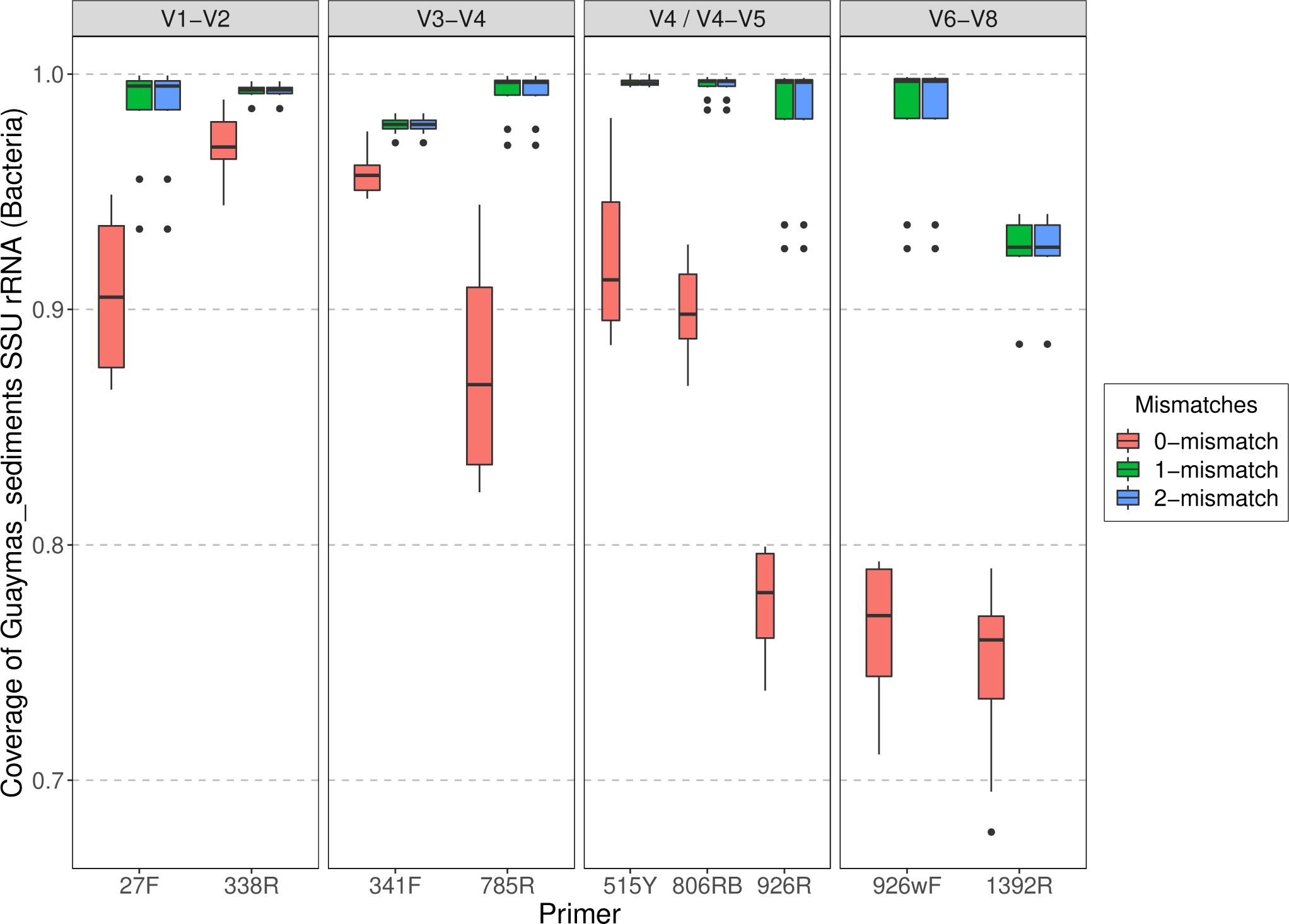
Predicted individual primer coverage allowing 0, 1, and 2-mismatches for *Bacteria* (not including *Cyanobacteria* and plastids) using metagenomes derived from Guaymas Basin sediment samples.

**Fig S16:**
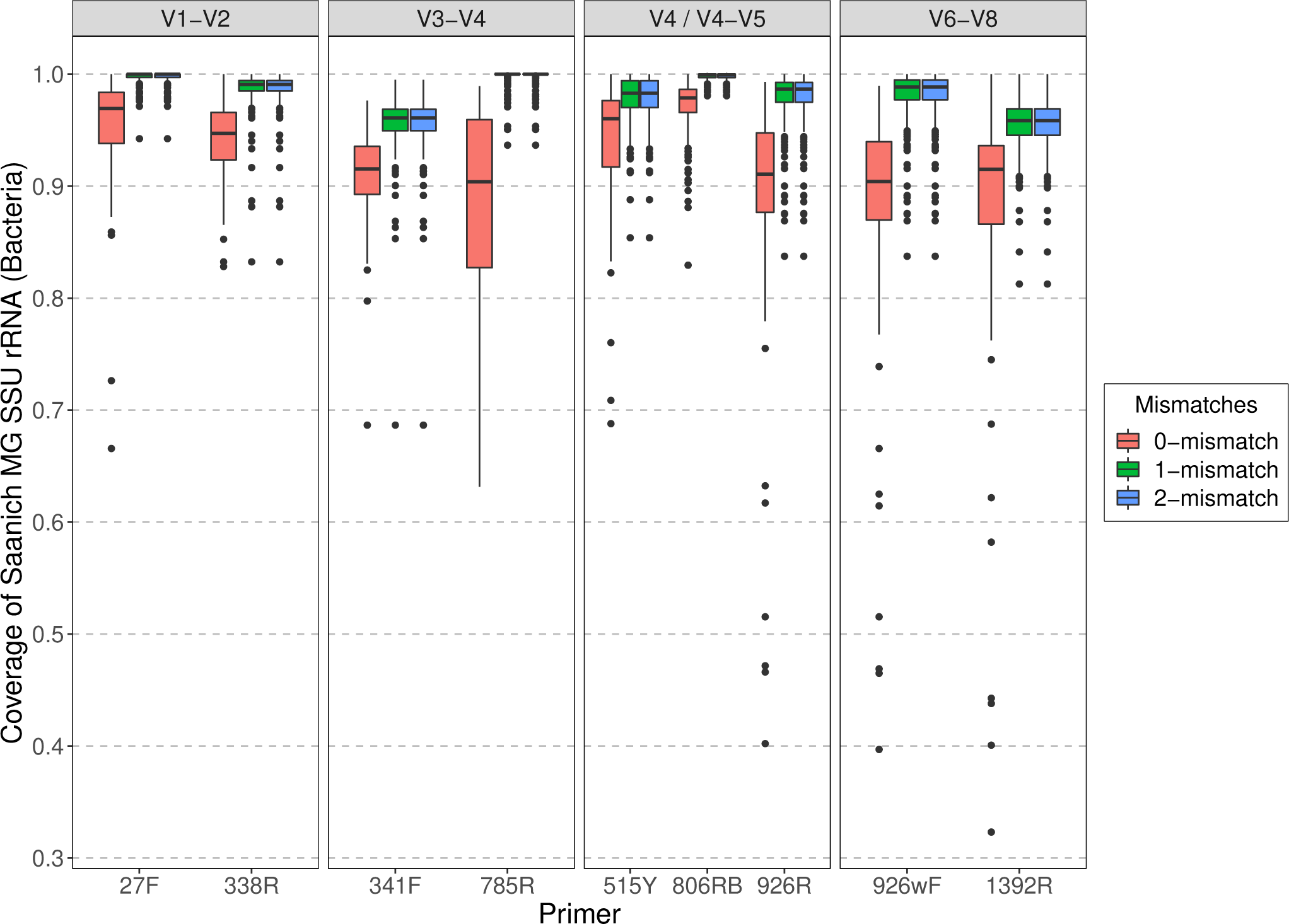
Predicted individual primer coverage allowing 0, 1, and 2-mismatches for *Bacteria* (not including *Cyanobacteria* and plastids) using metagenomes derived from Saanich Inlet water samples.

**Fig S17:**
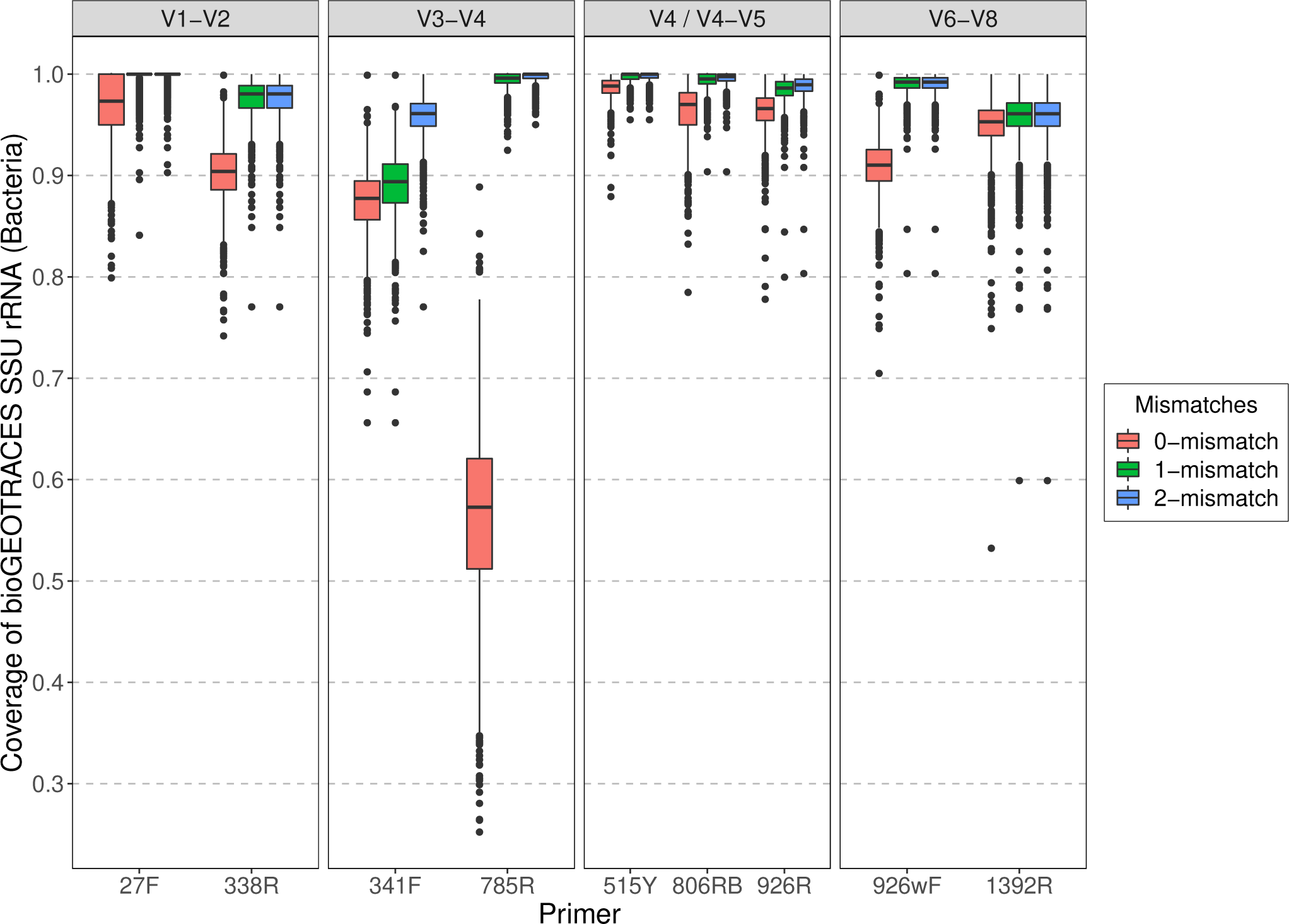
Predicted individual primer coverage allowing 0, 1, and 2-mismatches for *Bacteria* (not including *Cyanobacteria* and plastids) using metagenomes derived from BioGEOTRACES/BATS/HOT water samples.

**Fig S18:**
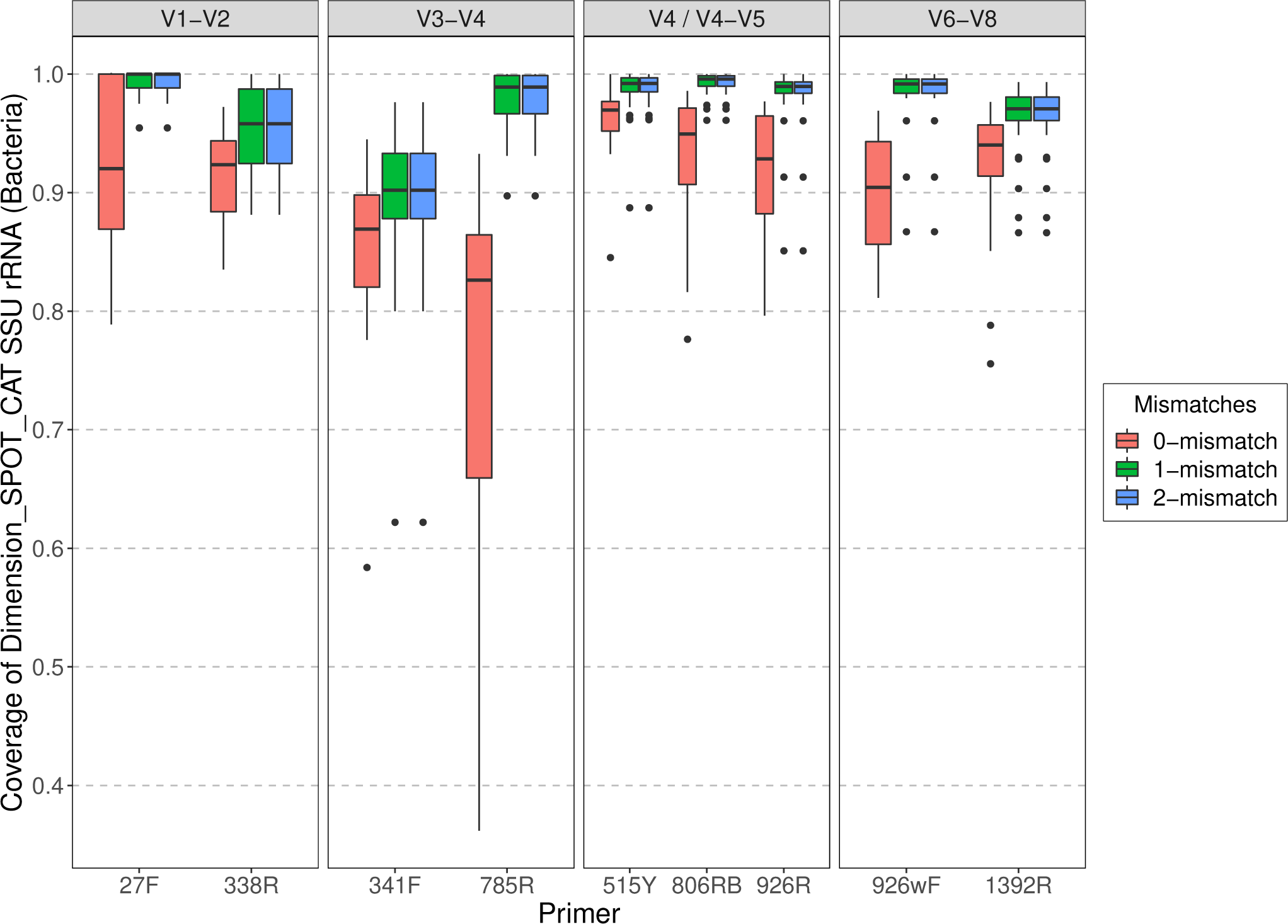
Predicted individual primer coverage allowing 0, 1, and 2-mismatches for *Bacteria* (not including *Cyanobacteria* and plastids) using metagenomes derived from SPOT/Catalina water samples.

**Fig S19:**
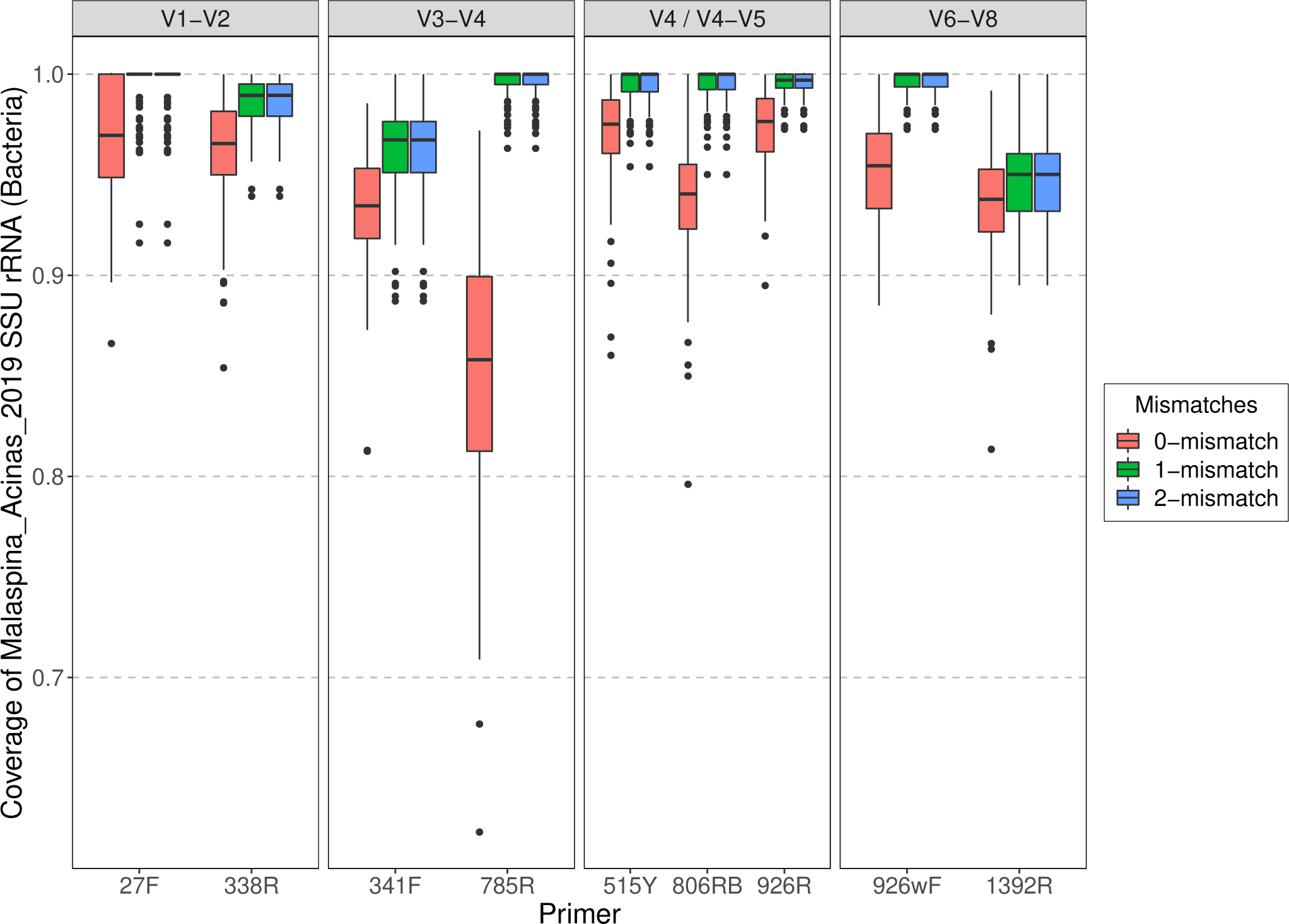
Predicted individual primer coverage allowing 0, 1, and 2-mismatches for *Bacteria* (not including *Cyanobacteria* and plastids) using metagenomes derived from Malaspina deep-sea water samples.

**Fig S20:**
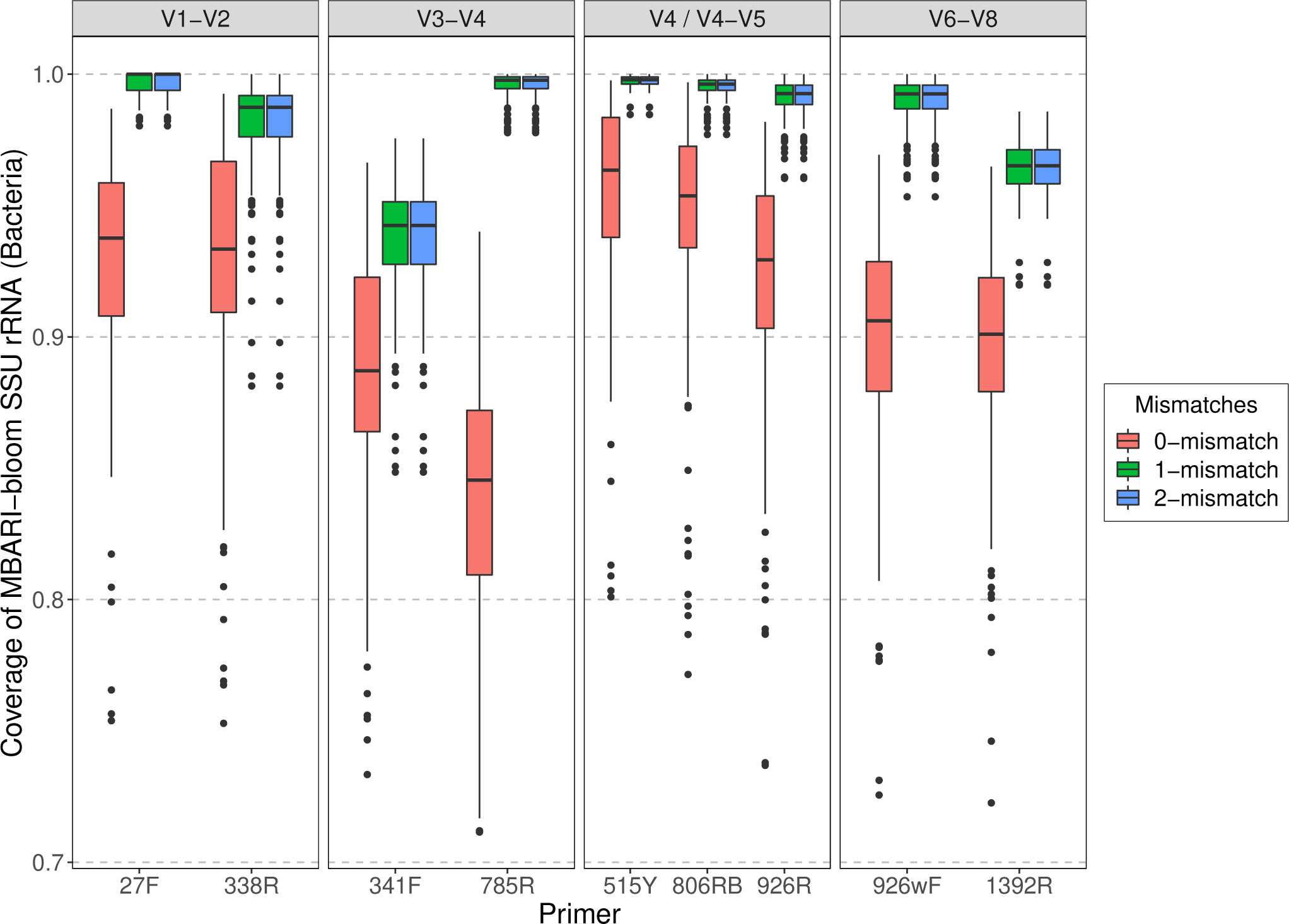
Predicted individual primer coverage allowing 0, 1, and 2-mismatches for *Bacteria* (not including *Cyanobacteria* and plastids) using metagenomes derived from MBARI bloom water samples.

**Fig S21:**
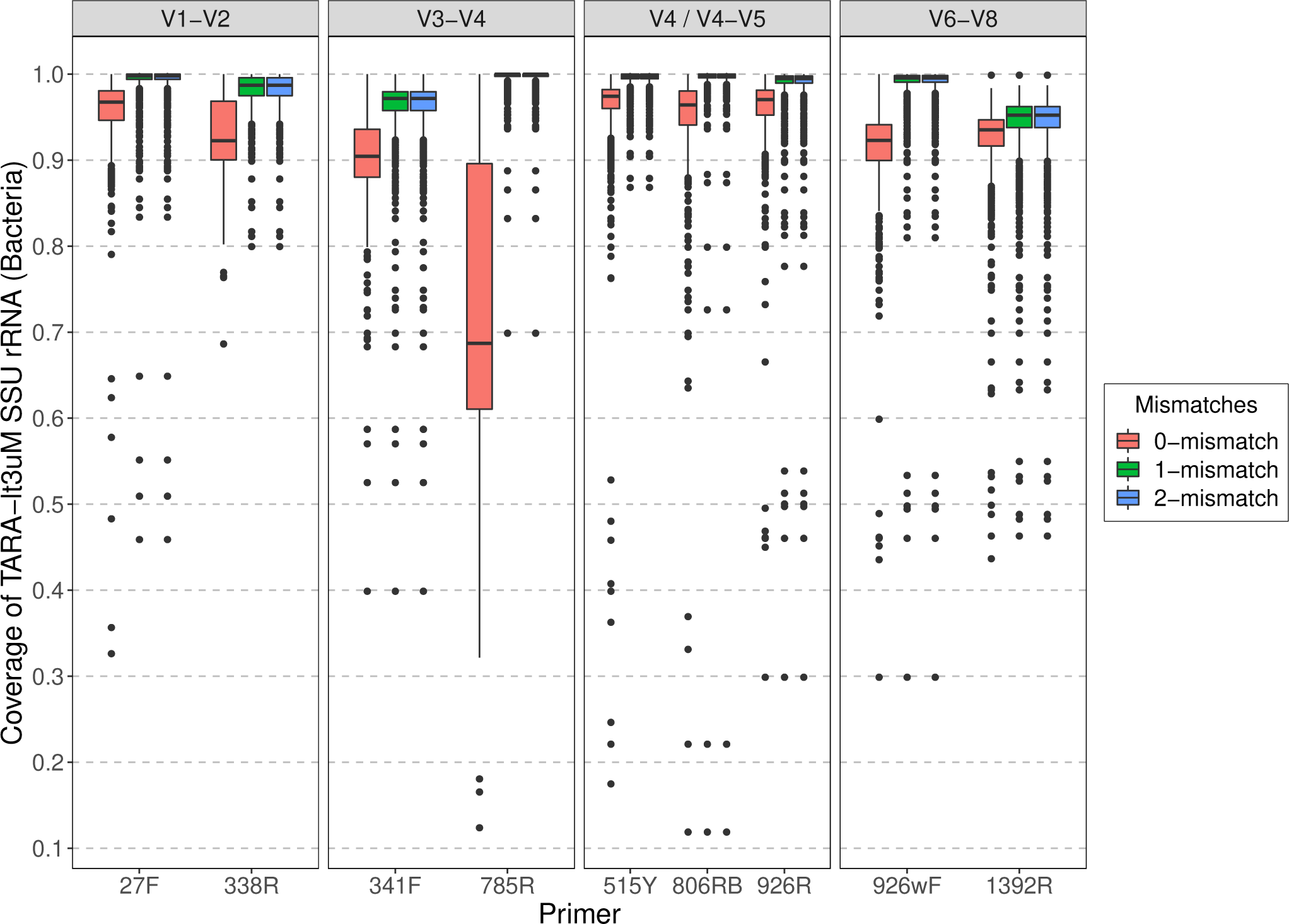
Predicted individual primer coverage allowing 0, 1, and 2-mismatches for *Bacteria* (not including *Cyanobacteria* and plastids) using metagenomes derived from Tara Oceans water samples (<3µm).

**Fig S22:**
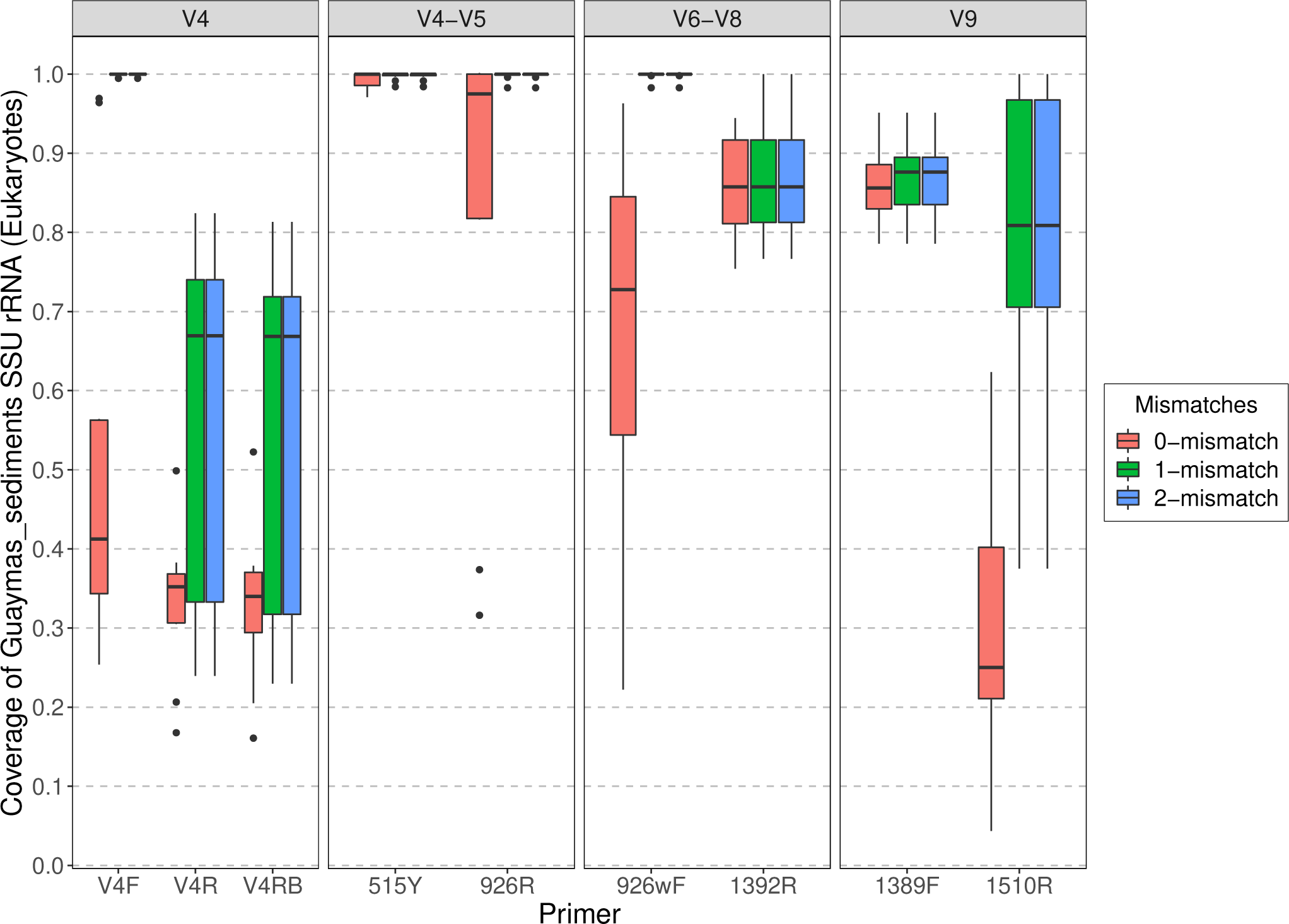
Predicted individual primer coverage allowing 0, 1, and 2-mismatches for *Eukarya* using metagenomes derived from Guaymas Basin sediment samples.

**Fig S23:**
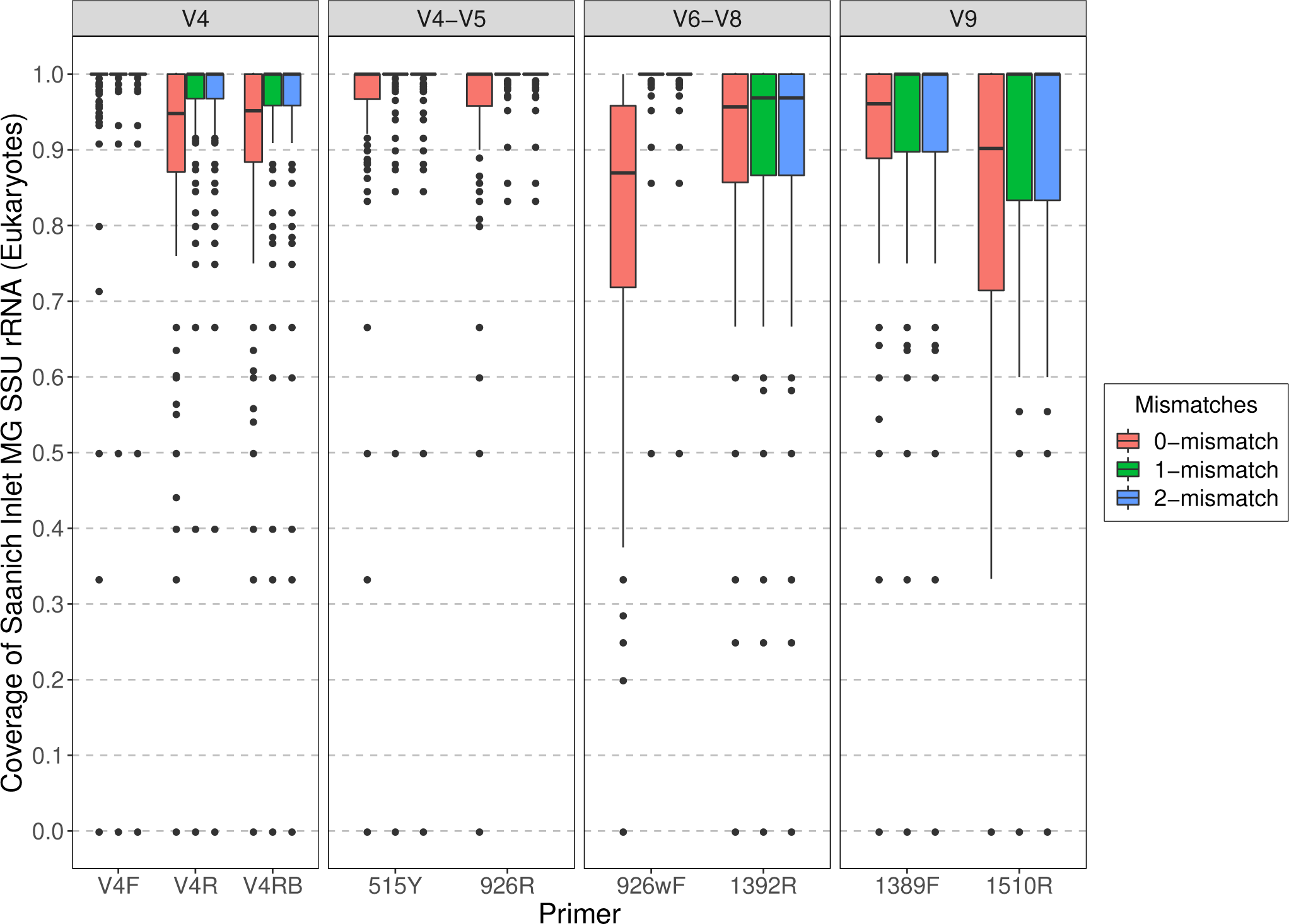
Predicted individual primer coverage allowing 0, 1, and 2-mismatches for *Eukarya* using metagenomes derived from Saanich Inlet water samples.

**Fig S24:**
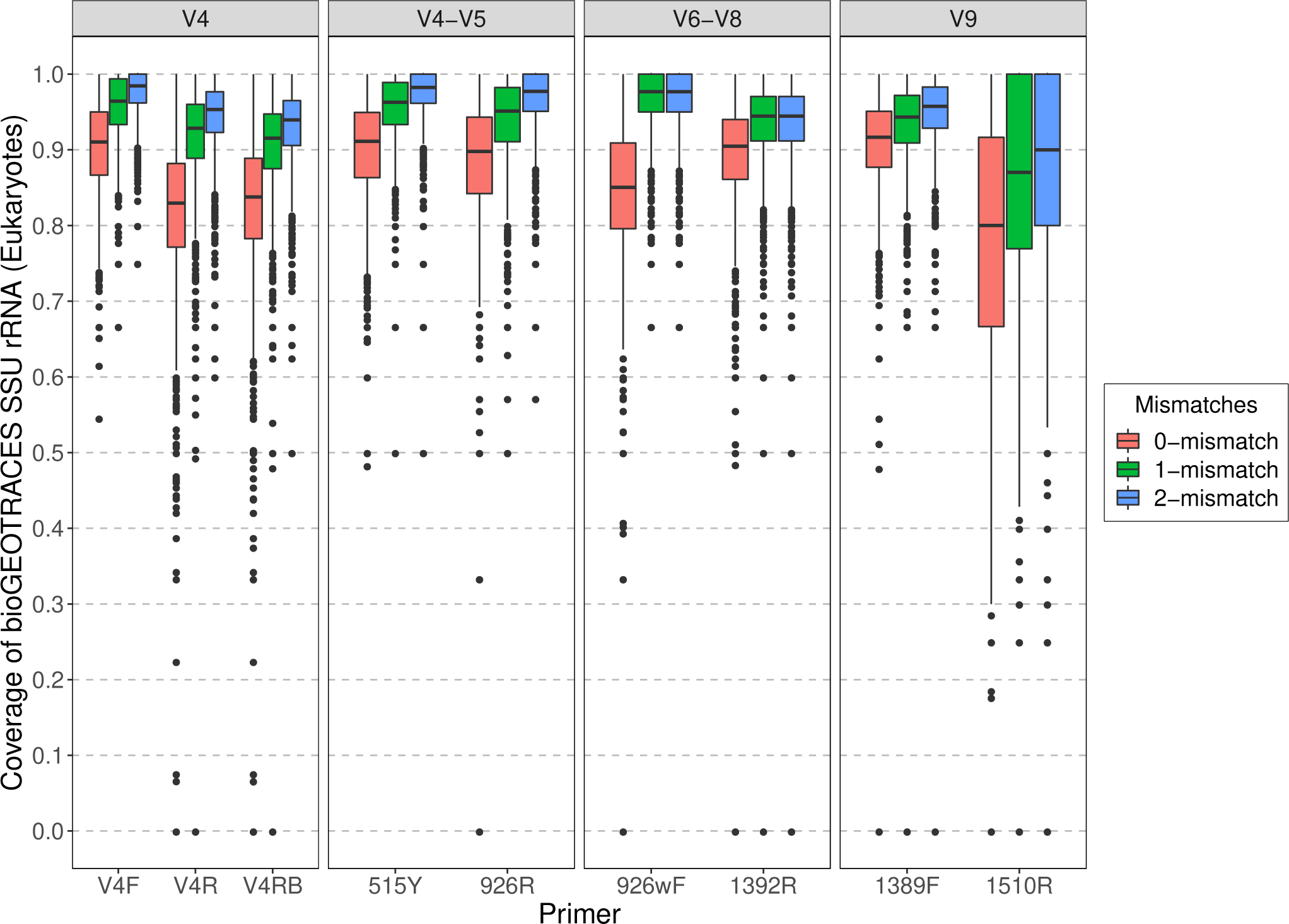
Predicted individual primer coverage allowing 0, 1, and 2-mismatches for *Eukarya* using metagenomes derived from BioGEOTRACES/BATS/HOT water samples.

**Fig S25:**
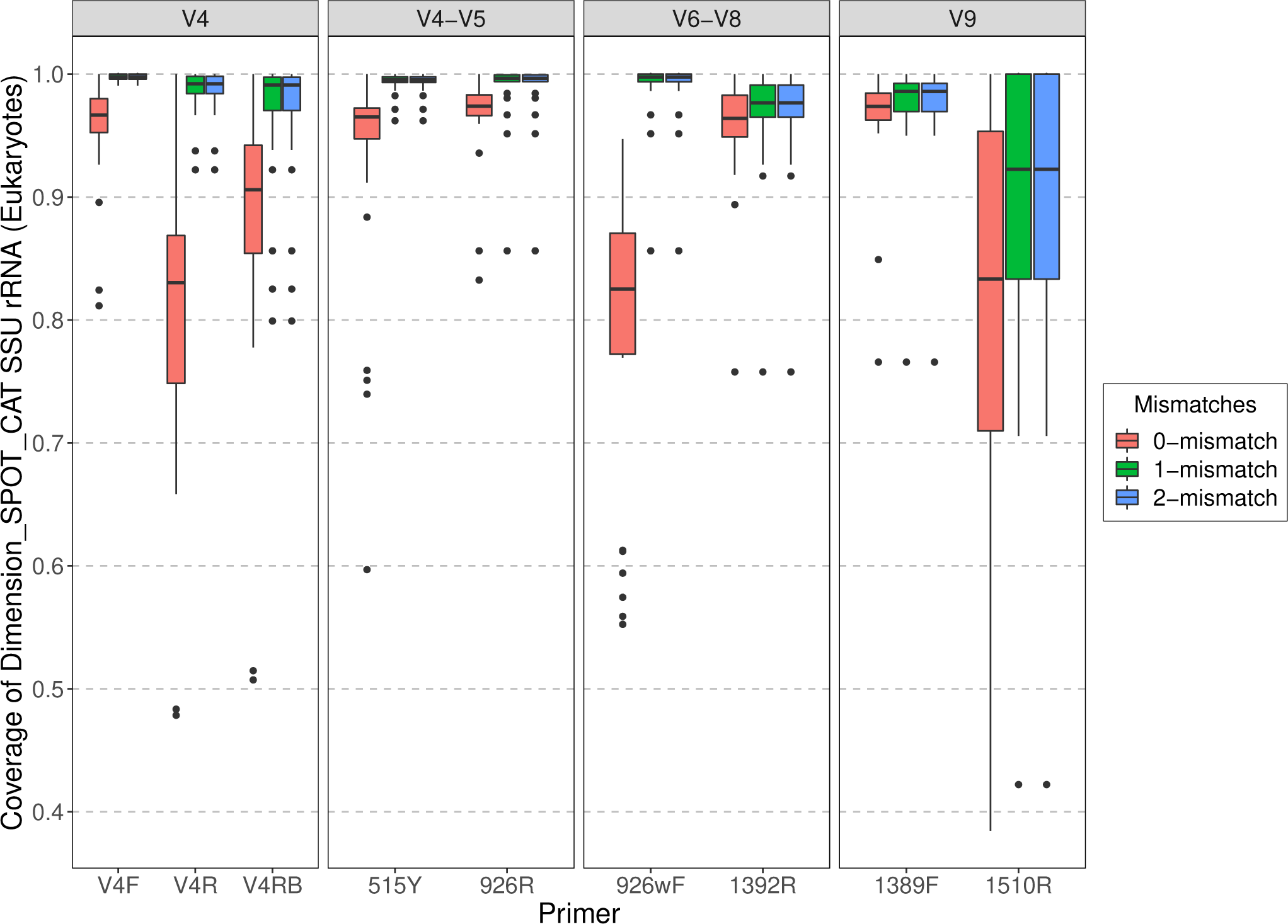
Predicted individual primer coverage allowing 0, 1, and 2-mismatches for *Eukarya* using metagenomes derived from SPOT/Catalina water samples.

**Fig S26:**
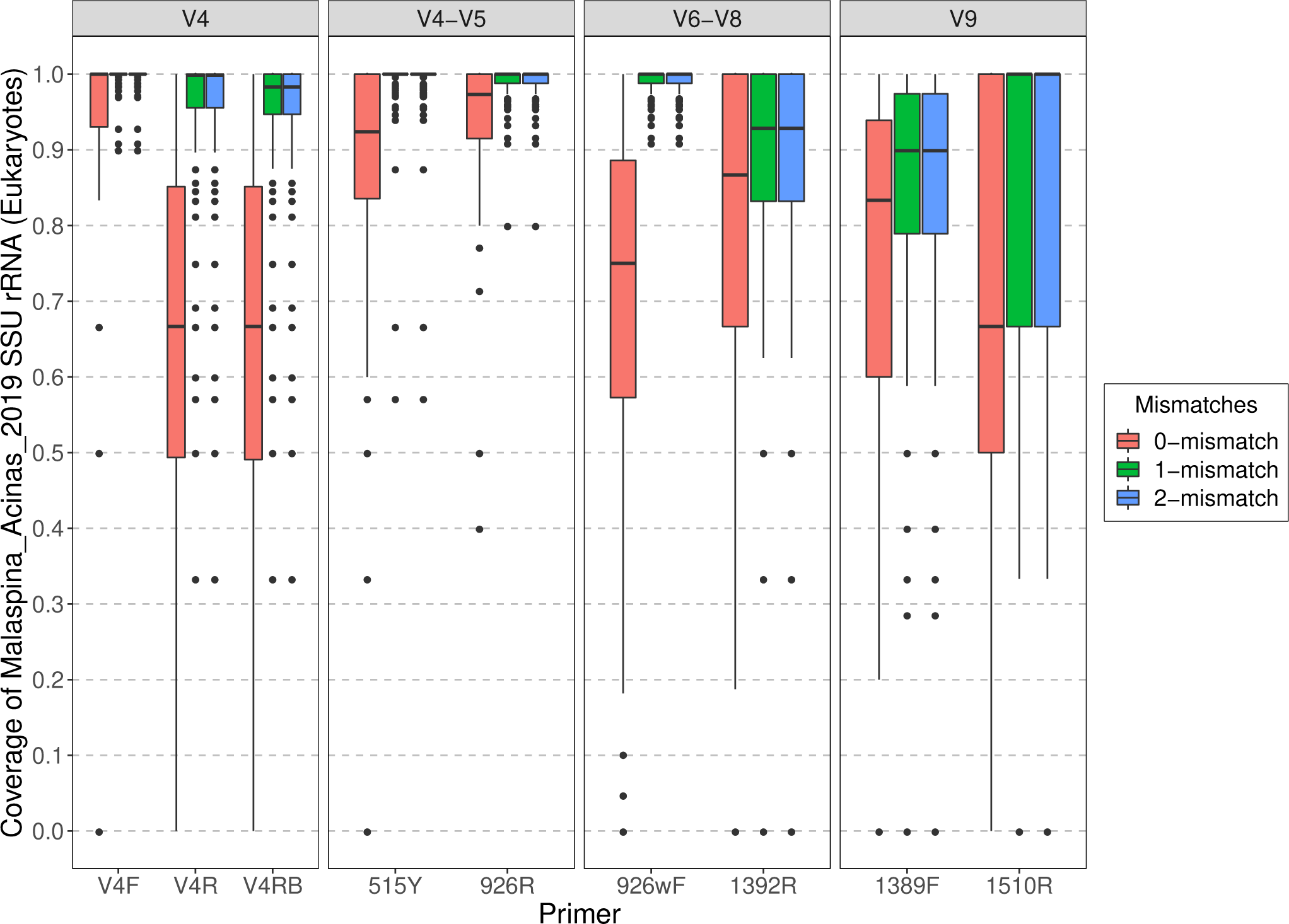
Predicted individual primer coverage allowing 0, 1, and 2-mismatches for *Eukarya* using metagenomes derived from Malaspina deep-sea water samples.

**Fig S27:**
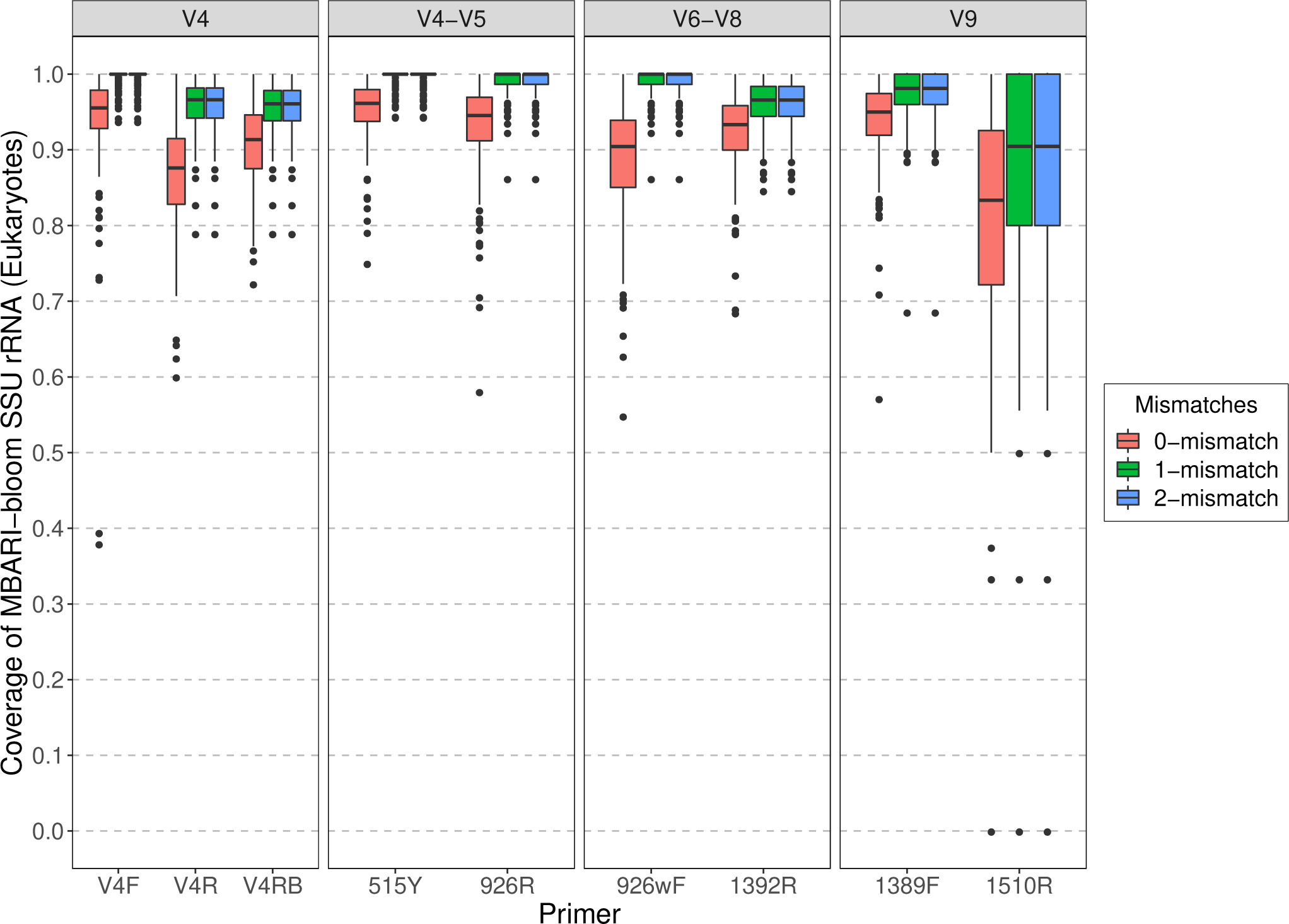
Predicted individual primer coverage allowing 0, 1, and 2-mismatches for *Eukarya* using metagenomes derived from MBARI bloom water samples.

**Fig S28:**
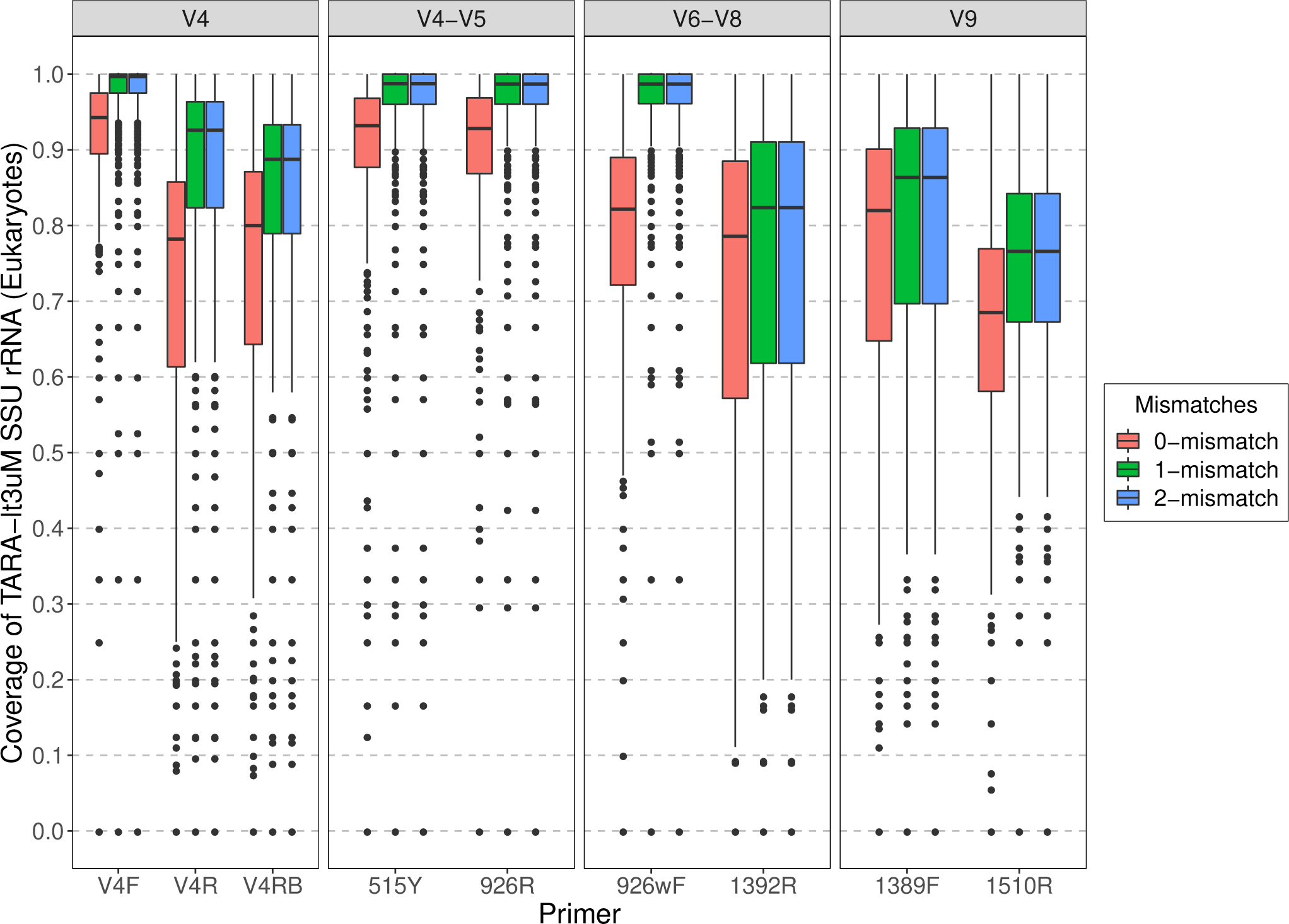
Predicted individual primer coverage allowing 0, 1, and 2-mismatches for *Eukarya* using metagenomes derived from Tara Oceans water samples (<3µm).

**Figure S29:**
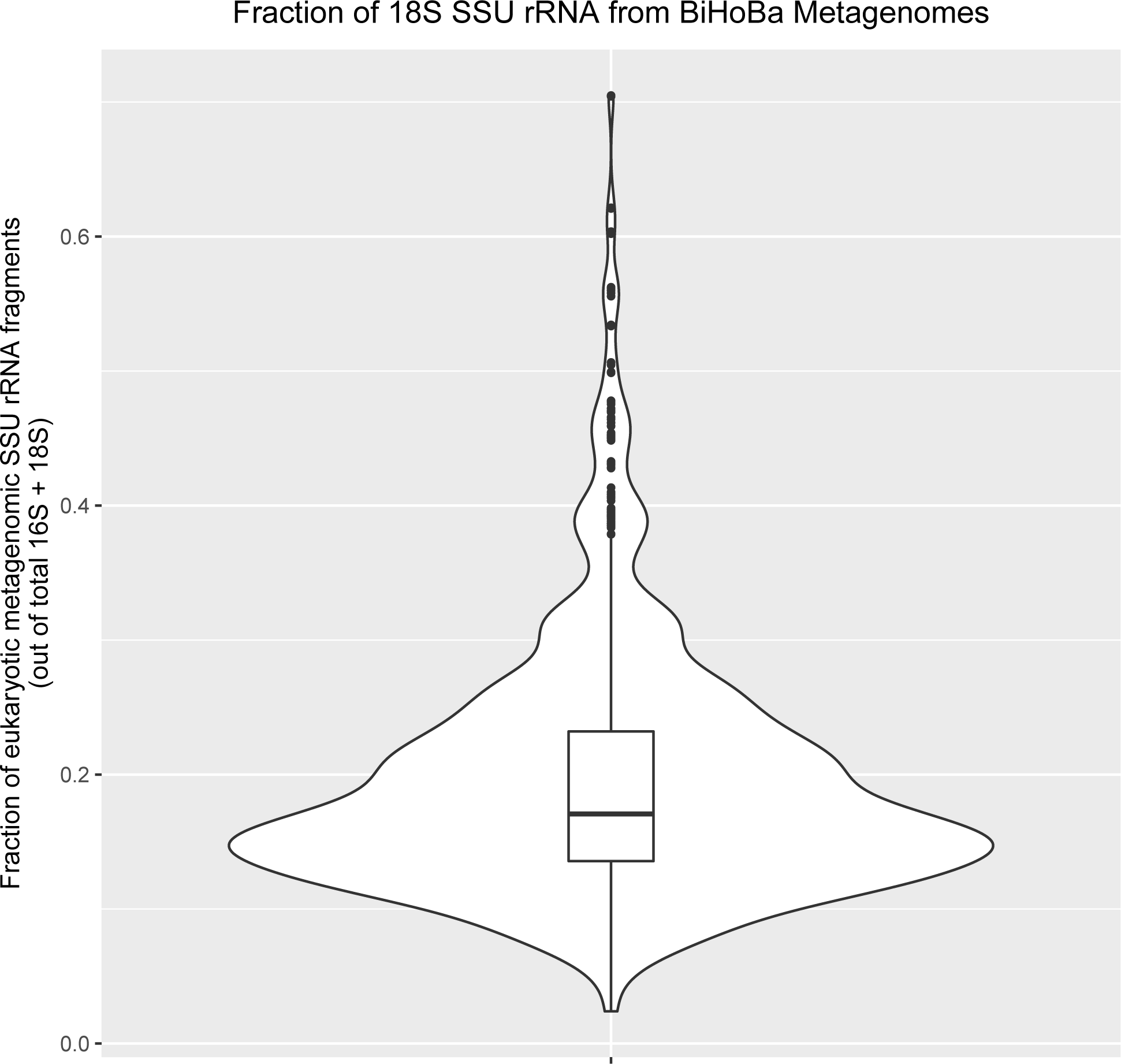
Violin plot with inlaid boxplot showing the fraction of eukaryotic (18S) SSU rRNA as a fraction of total across BATS, HOT, and bioGEOTRACES metagenomic datasets. The violin plot provides a visual description of the distribution of data - i.e. in this case showing that the vast majority of samples had < 30% of SSU rRNA reads accounted for by 18S rRNA sequences.

**Figure S30:**
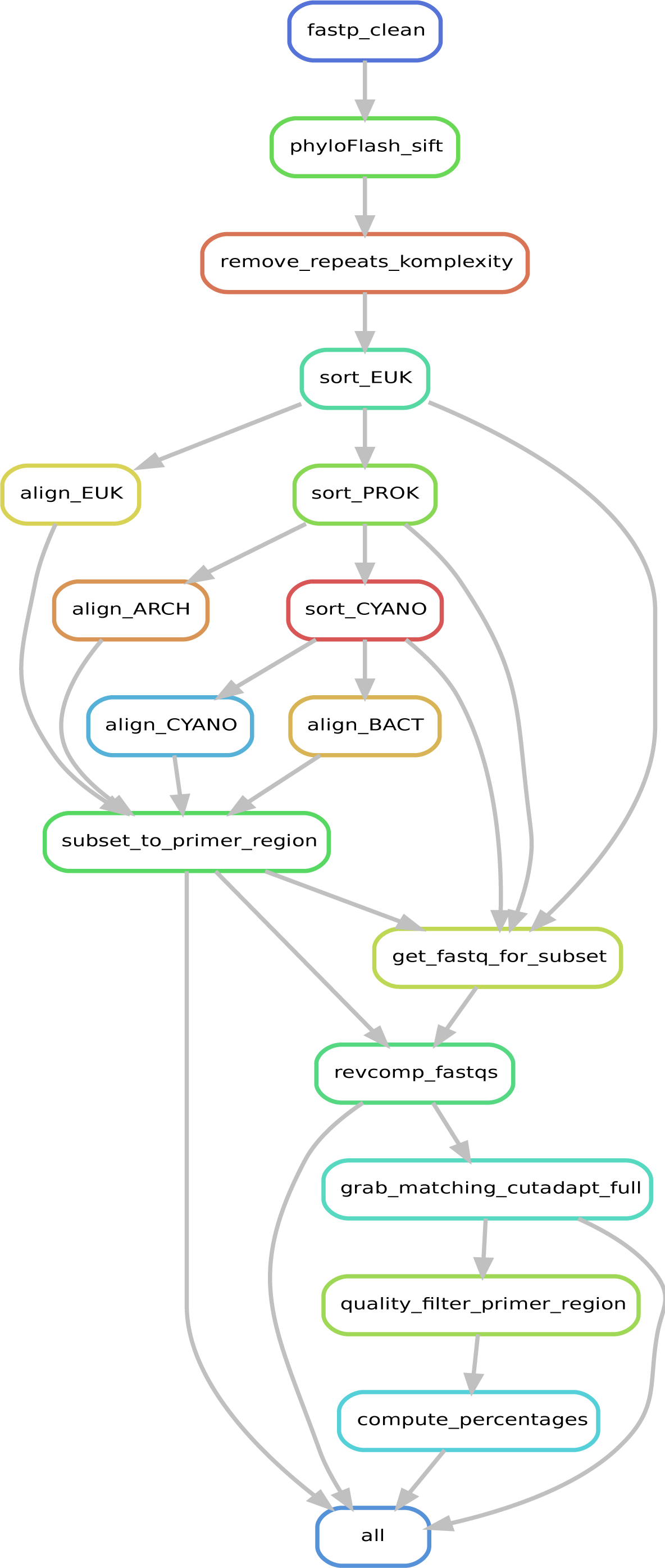
Rulegraph for “Snakefile- compute.smk” portion of workflow. Briefly, SSU rRNA is identified using phyloFlash from fastp quality-filtered fastq reads, which are then sorted with bbsplit.sh into 4 categories, and aligned to SSU rRNA references with pyNAST after removing low-complexity sequences. For each group and primer, matches to primer regions are then identified using custom python scripts and cutadapt (at 0, 1, 2, and 6- mismatches), quality filtered to omit sequences where the primer region plus 5 leading and trailing bases has one individual base with a phred score below 30. Matches are then summarized in plain-text output and ggplot2 graphs.

**Figure S31:**
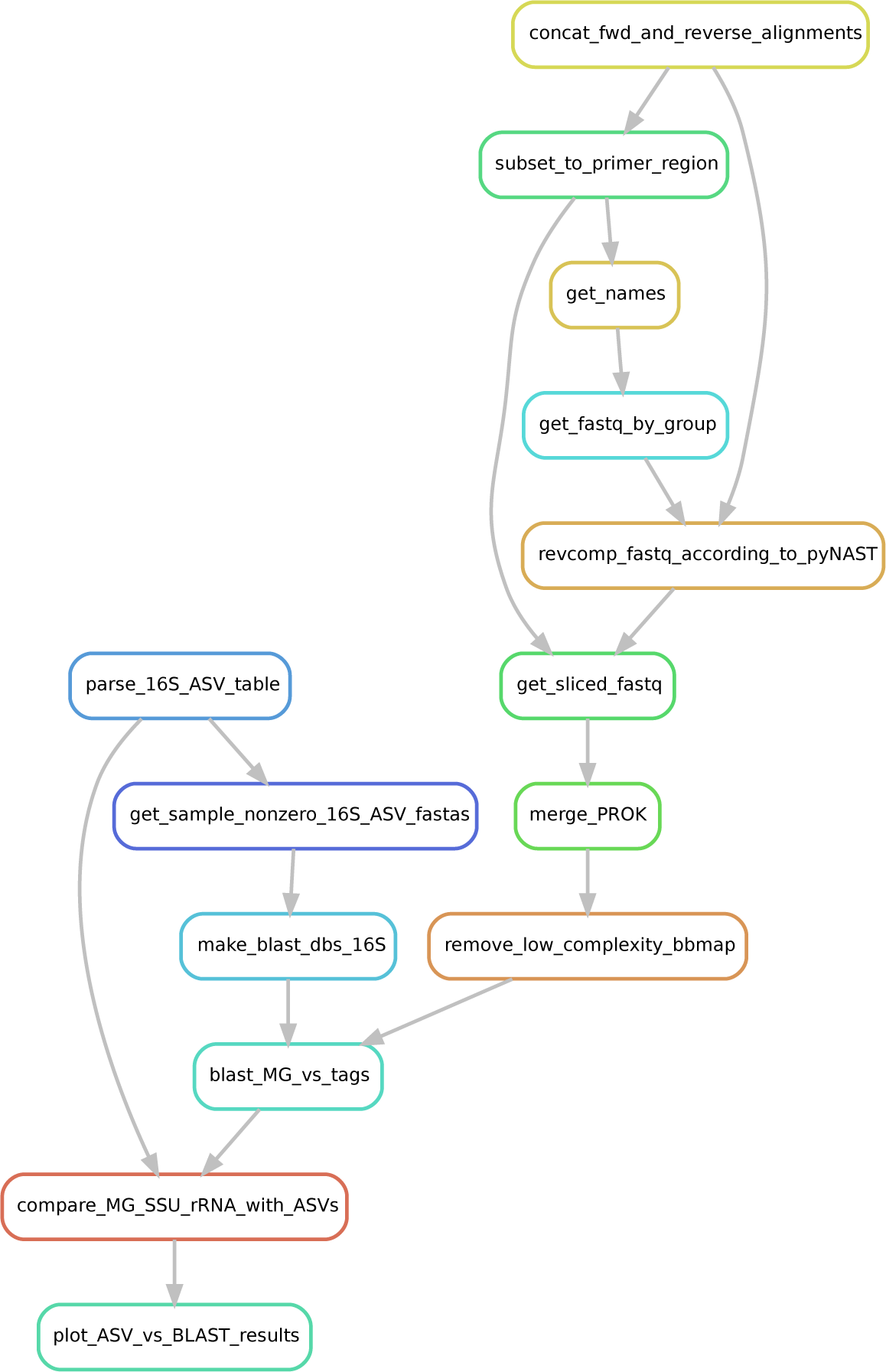
Rulegraph for “Snakefile- compare.smk” portion of workflow to intercompare metagenome and amplicon data.

